# Extensive genetic differentiation between recently evolved sympatric Arctic charr morphs

**DOI:** 10.1101/489104

**Authors:** Jóhannes Guðbrandsson, Kalina H. Kapralova, Sigríður R. Franzdóttir, Þóra Margrét Bergsveinsdóttir, Völundur Hafstað, Zophonías O. Jónsson, Sigurður S. Snorrason, Arnar Pálsson

## Abstract

The availability of diverse ecological niches can promote adaptation of trophic specializations and related traits, as has been repeatedly observed in evolutionary radiations of freshwater fish. The role of genetics, environment and history in ecologically driven divergence and adaptation, can be studied on adaptive radiations or populations showing ecological polymorphism. Salmonids, especially the *Salvelinus* genus, are renowned for both phenotypic diversity and polymorphism. Arctic charr (*Salvelinus alpinus*) invaded Icelandic streams during the glacial retreat (about 10,000 years ago) and exhibits many instances of sympatric polymorphism. Particularly well studied are the four morphs in Lake Þingvallavatn in Iceland. The small benthic (SB), large benthic (LB), planktivorous (PL) and piscivorous (PI) charr differ in many regards, including size, form and life history traits. To investigate relatedness and genomic differentiation between morphs we identified variable sites from RNA-sequencing data from three of those morphs, and verified 22 variants in population samples. The data reveal genetic differences between the morphs, with the two benthic morphs being more similar and the PL-charr more genetically different. The markers with high differentiation map to all linkage groups, suggesting ancient and pervasive genetic separation of these three morphs. Furthermore, GO analyses suggest differences in collagen metabolism, odontogenesis and sensory systems between PL-charr and the benthic morphs. Genotyping in population samples from all four morphs confirms the genetic separation and indicates that the PI-charr are less genetically distinct than the other three morphs. The genetic separation of the other three morphs indicates certain degree of reproductive isolation. The extent of gene flow between the morphs and the nature of reproductive barriers between them remain to be elucidated.

## 1 Introduction

Organismal diversity reflects the process of evolution and highlights the importance of natural selection in building and maintaining adaptations (Darwin 1859). While purifying selection preserves adaptations and biological functions, positive selection alters phenotypic traits and frequencies of genetic variations influencing them (Via 2001, 2009). Feeding is a primary function in all animals and in nature, we see many examples of spectacular adaptive radiation where natural selection has generated a range of adaptations in specific feeding structures and foraging behaviour (Losos and Ricklefs 2009; Seehausen and Wagner 2014). Recently, studies have demonstrated how positive selection can shift phenotypic distributions on generational time scales (Grant and Grant 2002; Nosil 2012). The recurrent ecological specializations, like for feeding (Seehausen and Wagner 2014), found in many species and populations of freshwater fish, represent interesting study systems and set of functional phenotypes for analyses of evolutionary change, convergence and parallel adaptation (Muschick et al. 2012).

### 1.1 Genome wide divergence or islands of differentiation?

The genomics revolution has spawned powerful tools for studying relatedness of populations and species, the connections between genetic and phenotypic variation, e.g. how variations in the form of feeding structures are caused by differential expression of key developmental genes (Abzhanov et al. 2006; Guðbrandsson et al. 2018), and to determine to what extent these differences are due to genetic differences (and how they are distributed in genomes) (Wolf and Ellegren 2016). These methods have enabled studies evaluating feedback between the organism and its environment, the role of plasticity in generating functional variation and possibly promoting adaptation (Morris and Rogers 2014; Abouheif et al. 2014). Central to this study, genomic methods can reveal differences between populations and ecotypes, and highlight genes and pathways that may be under positive selection in natural populations (Malinsky et al. 2015; Wolf and Ellegren 2016).

The differentiation and divergence between related groups (populations, sub-species or species) are influenced by genomic parameters, population history and evolutionary forces (Seehausen et al. 2014; Vijay et al. 2017). The genetic separation may be rather uniform over the entire genome or localized to “genomic islands of differentiation”, depending on multiple factors, e.g. the strength of selection, the nature of the adaptive traits, genomic structure, population genetic parameters and history (Malinsky et al. 2015; Wolf and Ellegren 2016). Studies of closely related groups can detect the genetic correlates of adaptive traits (Pease et al. 2016), but this is not straightforward for a number of reasons. Selection operates on multiple genes influencing all aspects of fitness, at the same time, for instance standing variation and recent variants (Vijay et al. 2017), alleles derived from introgression, even possibly balanced ancestral polymorphism (Pardo-Diaz et al. 2012; Guerrero and Hahn 2017). Furthermore, recently separated groups adapting to different environments can be viewed as being scattered along a “speciation continuum” (Theis et al. 2014; Seehausen et al. 2014). Under this view, changes in environmental or population genetic parameters can move a population along this continuum (Grant and Grant 2008). Specifically, such changes can lead to the merging of previously distinct populations, speed up their divergence or produce reproductively isolated species (Hendry et al. 2009; Seehausen et al. 2014; Lowry and Gould 2016). We are keen to explore the genetic differentiation of recently evolved populations within a species, that coexist in the same lake, with the aim to study the genetics of ecological specializations.

### 1.2 Resource polymorphism along a benthic - limnetic axis

It is widely acknowledged that varying resource utilization of allopatric populations can lead to divergence of traits which, given time, can result in adaptive divergence among these populations. However, discrete variation in resource use among individuals in the same area, e.g. foraging in different habitats, can also generate divergent traits within populations (Skúlason and Smith 1995; Bernatchez et al. 2010). The occurrence of discrete phenotypes (morphs) of a species living in sympatry and diverging in traits that relate to the utilization of different resources (food, breeding grounds, etc) has been called resource polymorphism. Typically, diverging traits of resource morphs involve morphology, behaviour and/or life history characteristics (Skúlason and Smith 1995; Smith and Skúlason 1996). Resource polymorphism can arise through developmental plasticity within a homogenic population, by natural selection on genetically encoded traits or a combination of these mechanisms. For instance, the broad and narrow-headed morphs of the European eel (*Anguilla anguilla*) were found to be determined to a large extent by diet (De Meyer et al. 2016) and ecomorphs of killer whales (*Orcinus orca*) were hypothesized to have originated through plastic responses of a small founder population (Foote et al. 2016). Examples of genetically determined resource polymorphism can be found in crater lake cichlid fishes, e.g. were a single locus effects jaw and body shape (Fruciano et al. 2016). Likewise, in benthic and limnetic “species” of threespine stickleback (*Gasterosteus aculeatus*), the same loci show signs of differentiation (Jones et al. 2012).

Salmonids are renowned for their phenotypic diversity, both among and within populations, with multiple examples of resource polymorphism. The most phenotypically diverse and polymorphic species seem to be in the *Salvelinus* genus, including Arctic charr *Salvelinus alpinus* (Klemetsen 2013), Lake charr (also called Lake trout) *Salvelinus namaycush* (Muir et al. 2016) and Dolly Varden charr (*Salvelinus malma*) (also called Dolly Varden trout) with as many as seven morphs found in the same lake (Markevich et al. 2018). Arctic charr colonized lakes and rivers on the northern hemisphere after the last glaciation period (approx 9,000-12,000 years ago) (Snorrason and Skúlason 2004; Noakes 2008; Klemetsen 2010). In Iceland, multiple lakes harbour polymorphic Arctic charr (Skúlason et al. 1992; Snorrason and Skúlason 2004; Wilson et al. 2004; Woods et al. 2012b) and a unique small benthic morphotype is found in many streams and ponds across the country, especially in cold springs with a lava-rock bottom in the geologically younger parts of the island (Kapralova et al. 2011; Kristjánsson et al. 2012). One of four sympatric charr morphs found in Lake Þingvallavatn, Iceland’s largest lake, is of this type. Population genetics show that populations of Arctic charr in Iceland group by geography, not morphotype (Gíslason 1998; Kapralova et al. 2011), supporting the notion that small benthic morphs and other derived forms have evolved independently in different locations. For instance, the four Lake Þingvallavatn morphs are more closely related to one another than to other charr populations (Kapralova et al. 2011). Genetic analysis also indicates closely related sympatric morphs in Norway, Scotland and Transbaikalia (Hindar et al. 1986; Gordeeva et al. 2015; Jacobs et al. 2018), while in other cases sympatric charr morphs seem to have emerged by more than one invasion (Verspoor et al. 2010).

### 1.3 The four sympatric charr morphs in Lake Þingvallavatn

Lake Þingvallavatn formed in a rift zone as the Icelandic ice-cap receded ∼10,000 years ago and was shaped by volcanic activity and the isostatic rebound. The lake has been influenced by tectonic movements, causing extensive subsidence and horizontal extension with extensive rift-forming in the central graben (Saemundsson 1992). The lake harbours four distinct Arctic charr morphs: Small benthic (SB), Large benthic (LB), Planktivorous (PL) and Piscivorous (PI) charr, that differ along a benthic - limnetic axis and this is reflected in their form, size, habitat use, diet, life history characteristics and parasite loads (Sandlund et al. 1987; Jonsson et al. 1988; Malmquist et al. 1992; Frandsen et al. 1989; Kapralova et al. 2013) and represent genuine resource polymorphism (Skúlason and Smith 1995) (Figure 1). The morphs also differ extensively in spawning times (Skúlason, Snorrason, Noakes, Ferguson and Malmquist 1989). Common garden experiments on embryos/offspring indicated heritable differences in various traits between morphs, e.g. morphology, sexual maturation rates, foraging behaviour (Skúlason, Noakes and Snorrason 1989; Skúlason et al. 1993, 1996; Kapralova et al. 2015) but plasticity was also found to be significant (Parsons et al. 2010, 2011).

**Figure 1:**
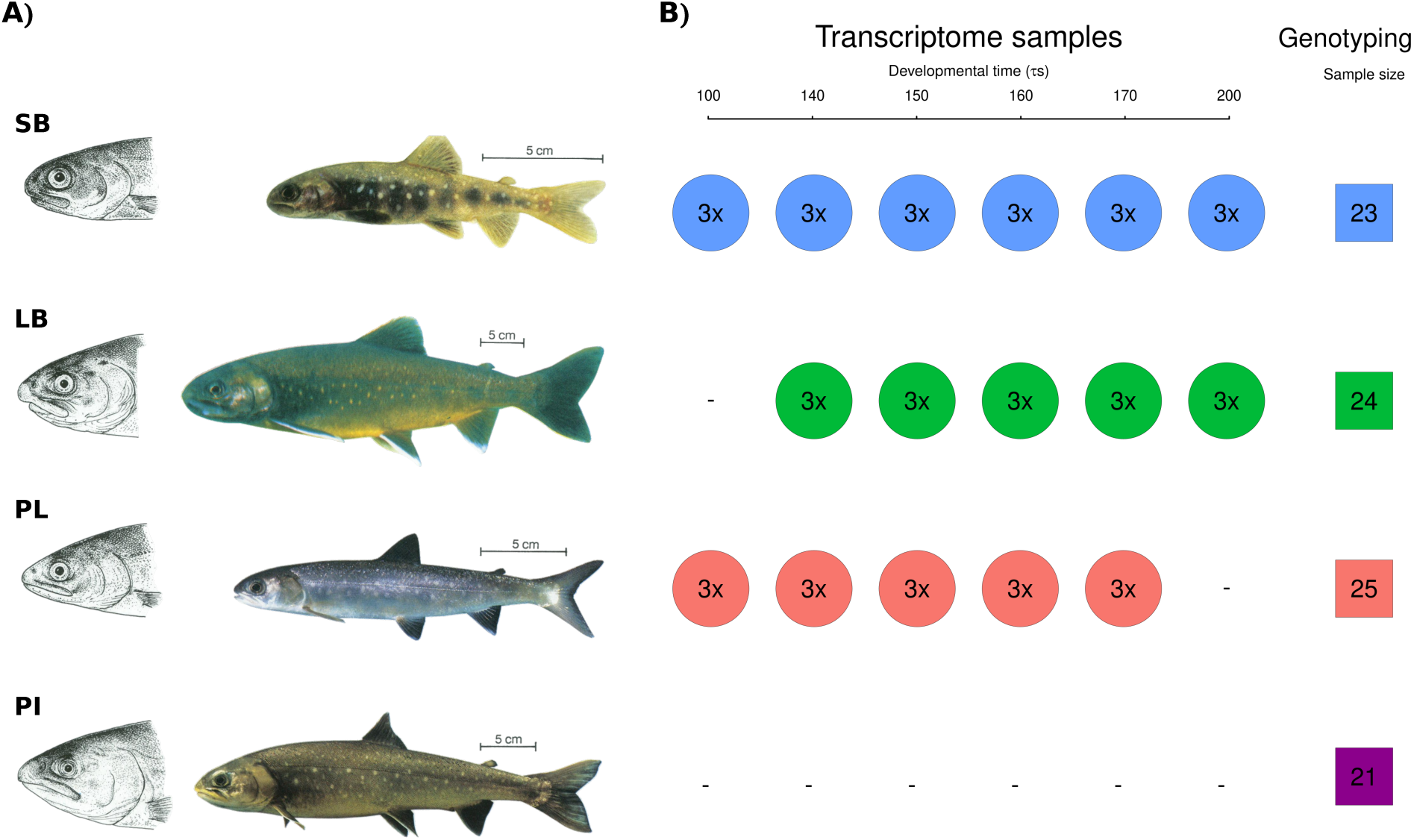
The phenotypically distinct sympatric Arctic charr from Lake Þingvallavatn and the sampling strategy. **A)** The four sympatric morphs are Small benthic (SB), Large benthic (LB), Planktivorous (PL) and Piscivorous (PI) charr. They differ in size (size bars = 5 cm), head and feeding morphology and pigmentation. Adapted from Sandlund et al. (1992) c Wiley-Blackwell, drawings by Eggert Pétursson. **B)** Sampling of charr for transcriptome sequencing (circles) and genotyping (squares) of population samples. The top 3 morphs were mined for genetic variation in the transcriptome and population samples were studied from all four, to confirm genetic variants. The transcriptome samples came from embryos at 6 developmental stages prior to hatching, from 100*τs* to 200*τs*, in the three morphs (circles)(Guðbrandsson et al. 2018). The sampling of each morph and developmental timepoint combination was replicated three times (biological replicates), each sample is a pool of mRNA from three embryos. Six timepoints were sampled of SB-charr, and five of LB- and PL-charr embryos. The population samples (squares) were obtained by gill netting on the spawning grounds, see methods. The morph colouring scheme (SB: blue, LB: green, PL: red and PI: purple) will be retained throughout the manuscript.

The earliest population genetic studies of the Lake Þingvallavatn charr found variation in some markers, but like in the RFLP analyses on mtDNA, generally weak if significant separation of morphs (Magnusson and Ferguson 1987; Danzmann et al. 1991; Volpe and Ferguson 1996). The estimated relationship of the morphs varied by studies (and markers), SB-charr (Magnusson and Ferguson 1987; Gíslason 1998), LB-charr (Kapralova 2008) and PL-charr (Volpe and Ferguson 1996) have all been estimated as the outgroup. PI-charr, on the other hand, tended clustered with both PL- and LB-charr. A study of 9 microsatellite markers in SB- and PL-charr from five spawning sites and LB-charr revealed significant but weak morph difference (average *F*_*ST*_ = 0.039) but did not resolve their relationship (Kapralova et al. 2011). Importantly, coalescence simulations on those data best supported a short initial phase of PL- and SB-charr allopatry followed by coexistence, rather than a pure sympatric origin of these morphs (Kapralova et al. 2011). More recent studies found loci with strong genetic differences (*F*_*ST*_ = 0.13 to 0.67) in two immunological genes (Kapralova et al. 2013) and rRNA genes within the mtDNA between morphs (Gudbrandsson et al. 2016).

Recent studies have studied the developmental and molecular correlates of morph divergence in the lake, by analyzing the morphology of embryos, miRNA and mRNA expression (Kapralova et al. 2014; Ahi et al. 2014; Kapralova et al. 2015; Gudbrandsson et al. 2016; Guðbrandsson et al. 2018). Most recently, RNA-sequencing of embryos of three of the four morphs (excluding PI-charr) reared in a common garden, revealed about 2000 transcripts with expression differences between morphs (Guðbrandsson et al. 2018). The PL-, LB- and SB-charr differed at the transcriptome level and the results suggested a closer relationship between the two benthic morphs (LB- and SB-charr), with PL-charr more distinct. Also, qPCR approaches have revealed differential expression between the morphs in the lake in genes related to extracellular matrix organization and skeletogenesis in early development (Ahi et al. 2013, 2014, 2015) and genes in the *mTOR*-pathway in muscles of SB-charr in Iceland (Macqueen et al. 2011).

Relatively few genomic resources have been available for Arctic charr, until recently when a reference genome was published (Christensen, Rondeau, Minkley, Leong, Nugent, Danzmann, Ferguson, Stadnik, Devlin, Muzzerall, Edwards, Davidson and Koop 2018). Reference genomes for other salmonid species have been made available in recent years, such as for Rainbow trout (*Oncorhynchus mykiss*, Berthelot et al. 2014; Pearse et al. 2018), Atlantic salmon (*Salmo salar*, Lien et al. 2016), Grayling (*Thymallus thymallus*, Varadharajan et al. 2018; Sävilammi et al. 2019), Chinhook salmon (*Oncorhynchus tshawytscha*, Christensen, Leong, Sakhrani, Biagi, Minkley, Withler, Rondeau, Koop and Devlin 2018) and one more (Coho salmon, *Oncorhynchus kisutch*) is available on NCBI (Genebank assembly GCA_002021735.1). About 88–103 million years ago an ancestor of Salmonids underwent a whole genome duplication (Ss4R), the fourth on the vertebrate lineage (Allendorf and Thorgaard 1984; Macqueen and Johnston 2014; Berthelot et al. 2014). Comparisons of several salmonids found significant synteny in their linkage groups (Danzmann et al. 2005; Sutherland et al. 2016), despite rearrangements (Timusk et al. 2011; Nugent et al. 2017). Thus a substantial fraction of genes in salmonids are paralogs in large syntenic regions (Nugent et al. 2017; Christensen, Rondeau, Minkley, Leong, Nugent, Danzmann, Ferguson, Stadnik, Devlin, Muzzerall, Edwards, Davidson and Koop 2018).

Here we mined the developmental transcriptome (Guðbrandsson et al. 2018) of three of the morphs (LB-, SB- and PL-charr) to test for genetic differences between them, elucidate their evolutionary relationship and identify loci and developmental pathways that may relate to morph differentiation. The data were used to evaluate three hypotheses about the causes of morph separation:

I. Because the salmonid’s homing behaviour leads offspring to spawn in the same location as their parents, heterogeneity in spawning places and micro-environments can lead to environmentally induced gene expression and phenotypic differences. In this scenario, morphs are environmentally induced, and genetic differences between them minor.
II. Ecological selection on specific traits has led to genetic separation of morphs, seen as allele frequency differences at variants in key genes related to fitness traits, in the face of gene flow between morphs (Seehausen et al. 2014; Wolf and Ellegren 2016). Under this model, genomic islands of differentiation are expected, with limited genetic differences between morphs in the rest of the genome.
III. Pre-zygotic barriers such as spatial and temporal separation in spawning between morphs (Skúlason, Snorrason, Noakes, Ferguson and Malmquist 1989) or behavioural/mating differences have kept the morphs reproductively isolated for some time (as has been indicated for SB- and PL-charr (Kapralova et al. 2011)). Under this scenario, modest genetic separation among the morphs is expected, on all linkage groups.

These three hypotheses are not mutually exclusive, and some may apply to specific pairs of morphs. The data enabled analyses of the genetic separation of three of the sympatric morphs, the origin of the rare piscivorous charr, the genomic patterns of differentiation between morphs and the genes and molecular systems that may associate with their specializations.

## 2 Materials and Methods

### 2.1 Sampling for transcriptome and population genetics

The fishing of parents for crosses from Lake Þingvallavatn and sampling of embryos for developmental transcriptome analysis is described in Guðbrandsson et al. (2018). Briefly, we caught SB-, LB- and PL-charr on the spawning grounds, ready to release eggs and milt, made crosses and reared embryos from each cross separately in a common garden at the Hólar Aquaculture station, Verið (Sauðárkrókur). Samples of embryos were taken at 5-6 developmental timepoints depending on morph (see Figure 1). The relative age in *τs*-units was defined be Gorodilov (1996) and is a non-linear function of time and temperature (see formula in Guðbrandsson et al. 2018). For each morph and developmental timepoint three biological replicates were analyzed, each of those isolated from pools of 3 embryos to a total of 48 samples (Figure 1). The crosses were made using multiple parents from the same morph. For most samples, the embryos came from crosses created in the 2010 spawning season (SB 150-200*τs*, PL 140-170 *τs*, LB 140-200*τs*). For SB- and PL-charr gametes from ten males and ten females were mixed for the cross but five parents of each sex for LB. Due to poor RNA quality of some samples, we had to add samples collected in 2011. For PL timepoint 100*τs* we used a cross from the 2011 spawning season with a pool of gametes, from 10 males and 10 females. For SB timepoints 100*τs* and 140*τs* we had to rely on two single pair crosses (one male and one female) due to difficulties of finding running SB-charr. Samples SB100A and SB100B came from one cross (referred to as SB-2011_A) and SB100C and the three samples for timepoint 140*τs* came from another (cross SB-2011_B).

For verification of single nucleotide polymorphism’s (SNP’s) and comparisons between morphs we sampled sexually mature individuals at the spawning grounds of the three morphs; 23 SB-, 24 LB- and 25 PL-charr, by gillnet fishing in Ólafsdrattur and Mjóanes in 2015. To gain insight the genetic status of the PI-charr we genotype 21 individuals which have been collect as by-catch in other surveys in the period 2010–2016. They could not be collected in a single season as PI-charr is the rarest morph and spawning grounds not as well defined as for the other morphs.

As before (Guðbrandsson et al. 2018), all fishing in Lake Þingvallavatn was with permissions obtained from the farmers in Mjóanes and from the Þingvellir National Park Commission. Ethics committee approval is not needed for regular or scientific fishing in Iceland (The Icelandic law on Animal protection, Law 15/1994, last updated with Law 55/2013).

### 2.2 RNA isolation and sequencing

Total mRNA was isolated from a pool of three embryos for each sample. The RNA was quality checked and used as a template for 48 cDNA libraries subjected to Illumina sequencing. The sequencing was performed on Hiseq2000 at deCODE genetics (Reykjavík, Iceland) yielding 1,188 million 101 paired-end reads. The sequencing reads from the 48 samples were deposited into the NCBI SRA archive under BioProject identifier PRJNA391695 and with accession numbers: SRS2316381 to SRS2316428. For a more detailed description about RNA isolation, quality checks and library construction see Guðbrandsson et al. (2018).

### 2.3 Transcriptome assembly, Abundance estimation and Annotation

The transcriptome assembly and annotation were described previously (Guðbrandsson et al. 2018), but briefly, the reads were quality trimmed and adapters removed using Trim Galore! (version 0.3.3, Krueger 2012). The filtered reads from all samples were assembled with Trinity (version v2.1.0, Grabherr et al. 2011). We used kallisto (version v0.42.4, Bray et al. 2016) to estimate transcripts abundance. To speed up the annotation process transcripts with fewer than 200 mapped reads were not retained for annotation, another reason being that lowly expressed transcripts are unlikely to supply variants with enough coverage to surpass quality filters. The transcripts were annotated with the Trinotate pipeline (version 2.0.2, Haas 2015). Orthologs of the transcripts in salmon (*Salmo salar*) and rainbow trout (*Oncorhynchus mykiss*) mRNA and protein sequences were found using `blastn` and `blastx` respectively (Altschul et al. 1990). Annotation from NCBI *Salmo salar* Annotation Release 100 (https://www.ncbi.nlm.nih.gov/genome/annotation_euk/Salmo_salar/100/, retrieved 2015-12-17) and SalmoBase (Samy et al. 2017, http://salmobase.org/, version from 2015-09-18) were both used for salmon and annotation for rainbow trout came from Berthelot et al. (2014, http://www.genoscope.cns.fr/trout/data/, version from 2014-05-19). Only the best match was retained for each reference database. For further details about the annotation process and parameters, we refer to Guðbrandsson et al. (2018).

### 2.4 Variant calling and quality filtering

To identify genetic variation in the transcriptome we mapped the quality-trimmed reads to the complete Trinity assembly (annotated and non-annotated transcripts). Pseudobam files generated by kallisto were supplied to eXpress (version 1.5.1 Roberts and Pachter 2012) to get single alignment for multi-mapping reads. eXpress uses expectation–maximization (EM) algorithm to get posterior probabilities for read placements and assigns reads to transcripts by the posterior probability. We used the default eXpress parameters except we set the batch option to 10 to get more EM-rounds and better assignment of reads. Reads with more than 10 mismatches were removed using samtools (version 1.1, Li et al. 2009) and bamtools (version 2.3.0, Barnett et al. 2011).

Candidate variants (hereafter variants) were called with FreeBayes (version v1.0.1-2-g0cb2697, Garrison and Marth 2012) on all the samples simultaneously. The coverage threshold was set to 480 reads and the threshold for a variant allele to be called to 48 reads. Due to our experimental setup of pooled sampling of three individuals we added the options `--pooled-discrete` and `--ploidy 6`. As we were sequencing RNA and to reduce run time other options used for FreeBayes were; `--use-duplicate-reads` and `--use-best-n-alleles 4`. Only bi-allelic variants were processed further. Variants were filtered based on coverage and allele frequency. We only consider positions with a minimum coverage of 10 reads in at least 30 samples, thereof the minimum of eight samples in each morph. Minimum allele frequency of the alternative allele was set to 10% and the maximum to 70%.

Due to the genome duplication in the salmonid ancestor (Allendorf and Thorgaard 1984; Moghadam et al. 2011) some of the candidate variants might reflect sequence divergence of paralogous genes rather than true segregating variants. We removed such candidates by identifying variants with negligible differences in allele frequency between samples (and thus individuals). Recall, each sample represents 3 individuals, with 6 copies of each chromosome. Thus, true biallelic segregating markers only have seven possible genotype combinations (0:6, 1:5, 2:4, 3:3, etc). This should result in allele frequency differences among samples, as it is highly unlikely that the same combination of genotypes will be found in all cases. In contrast, candidate variants due to fixed differences in paralogous genes will have similar allele frequency in all samples. To estimate this we calculated hierarchical F-statistics (Hartl and Clark 2006, p. 280-283) in R (R Core Team 2015). First, we estimated the variation between samples, with a statistic termed *F*_*PT*_; were *P* stands for pool (each sample is a pool of three individuals) and *T* the total population. We also calculated a traditional *F*_*ST*_, to capture variation in allele frequencies between morphs (subpopulations). Note, because of the sampling of related individuals the values of *F*_*ST*_ are not comparable to other studies (see results).

The *F*_*PT*_ statistic is analogous to the classical *F*_*IT*_, so low *F*_*PT*_ values indicate a low variation between samples just as low *F*_*IT*_ values indicate low variation among individuals (excess heterozygosity). Filtering on *F*_*PT*_ is therefore similar to removing markers that deviate from Hardy-Weinberg equilibrium when genotypes are on an individual basis. To determine a reasonable cutoff for *F*_*PT*_ we simulated 288 chromosomes with eight different alternative allele frequencies (0.01, 0.1, 0.2, 0.3, 0.4, 0.5, 0.6, 0.7). Chromosomes were distributed randomly among samples with the same experimental setup as in our study and hierarchical F-statistics were calculated. For each alternative allele frequency, 10,000 replicates were simulated. Based on this simulation (see figure S1) we chose *F*_*PT*_ = 0.1 as a cutoff and all variants below this value were removed. Samples with coverage of five reads or less were not included in the calculations of F-statistics.

We ran FreeBayes again on the filtered variants in order to phase fully coupled variants into longer haplotypes. We used a haplotype length window of 70 basepairs (bp, option `--haplotype-length 70`) and only asked for biallelic variants (`--use-best-n-allels 2`). Nevertheless, nine variants in the output were tri-allelic and therefore removed. We also removed variants if the new estimate of *F*_*PT*_ was the below the cutoff (0.1) in the new run. No further filtering of variants was performed.

### 2.5 Analysis of variant functionality and distribution among morphs

Open reading frame prediction with Transdecoder (Haas and Papanicolaou 2015) in the Trinotate pipeline (see above) was used to estimate the position (UTR, CDS) of the variants and determine if they affected the protein sequence. Further statistical analysis was performed in the R environment (R Core Team 2015) using the VariantAnnotation package (Obenchain et al. 2014) to handle the variant data. The alternative allele frequency (based on read counts) for each sample was used for further analysis of genetic separation between groups. To study the distribution of variation among samples we did principal component analysis (PCA) with the built in prcomp-function in R. Missing values in the dataset were populated with the mean alternative allele frequency for the variant (21087 out of 1848192 values were missing or 1.14%), prior to the PCA analyses. We calculated the mean allele frequency of each variant for each morph, and also the deviation from the other two as the sum of the allele frequency difference between them. For example the deviation for PL is *d*_*PL*_ = *d*(*PL, SB*) + *d*(*PL, LB*) where *d*(*PL, SB*) is the difference in mean allele frequency for PL- and SB-charr. To screen for morph separation, we calculated *F*_*ST*_ comparing allele frequency by morphs (see above). Variants with *F*_*ST*_ above 0.2 were analyzed for gene ontology (GO) enrichment. The GO tests were performed with the goseq-package in R (Young et al. 2010), separately for variants with the largest deviation of mean allele frequency for each morph. The goseq-package accounts for transcript length as more variants are expected in longer transcripts. All transcripts with variants (8,961 transcripts) were used as the reference set for GO-enrichment tests. We also ran GO tests on the gene level using annotation to Salmobase. Gene length was not taken into account in that case and the reference set consisted of 7317 genes. GO categories were also mapped to their ancestor using the GO.db-package in R (Carlson 2015). We only tested biological processes, omitting categories closer than 3 steps from the root of the annotation tree. The gene ontology annotation was based on SalmoBase, see above and Guðbrandsson et al. (2018). Categories with false discovery rate (Benjamini and Hochberg 1995) below 0.01 were considered significant. We used 0.01 instead of the classic 0.05 as six tests were conducted (each morph on transcript and gene level). We cataloged variants private to one morph, the criterion being less than 1% frequency of the alternative allele in the other morphs. Morph specific lists of private alleles were analyzed for GO enrichment the same way.

### 2.6 Genomic distribution of candidate variants

With the Arctic charr reference genome (Christensen, Rondeau, Minkley, Leong, Nugent, Danzmann, Ferguson, Stadnik, Devlin, Muzzerall, Edwards, Davidson and Koop 2018, Assembly `GCA_002910315.2 ASM291031v2`) it became possible to assign variants to linkage groups. A sequence 200 bp upstream and downstream of each variant in the transcriptome contigs were mapped to the genome with `blastn` within R (Hahsler and Nagar 2017) using: `-max_target_seqs 2 -max_hsps 12 -culling_limit 2`. In the case of more than one blast hit for each sub-sequence, hits with the highest bit score that included the variant was chosen. If no hit included the variant the hit with highest bit score was used. If more than one hit was equally likely, hits to chromosomes were priorities to hits to contigs or scaffolds. If equally likely hits mapped to the same chromosome or scaffold in the genome and were within 50kb from each other the hit with the lower position was chosen. The remaining variants were left unplaced.

### 2.7 Genotyping of candidate variants in a population sample

We genotyped 93 adult fish from the four morphs, PI- (21) PL- (24), LB- (24) and SB-charr (24). DNA was extracted with phenol chloroform according to standard protocols. Markers for genotyping were chosen based on high *F*_*ST*_ values, predicted biological functions of the genes they affected and for some to evaluate LD between linked markers. To design primers for genotyping we aligned regions around candidate variants, contigs from the assembly and similar regions in *O. mykiss* and *S. salar* (retrieved by `blast`). Locations where updated when the *S. alpinus* genome became available (Christensen, Rondeau, Minkley, Leong, Nugent, Danzmann, Ferguson, Stadnik, Devlin, Muzzerall, Edwards, Davidson and Koop 2018). Markers were also chosen to tag independent linkage groups, but a handful was picked to survey variation in specific chromosomal regions. Sequences surrounding 23 variants (Table S5) were submitted to LGC genomics which designed KASP TM primers (He et al. 2014). The reactions were run on an Applied Biosystems 7500 Real-Time PCR machine, with the standard KASP program. All but one set of primers (*mrpl52_T76A*) passed test runs and were run on 93 samples (with 3 blank controls). Analyses of genotype and allele frequencies, correspondence of genotypes to Hardy Weinberg proportions, and the correlation of allele frequencies in the transcriptome and charr populations (PL, LB and SB-charr) were conducted with R packages (`pegas, adegenet` and `hierfstat`) and custom made scripts (Jombart 2008; Paradis 2010; Goudet 2005).

## 3 Results

### 3.1 Genetic variation separating three sympatric Arctic charr morphs

To estimate the relatedness of sympatric charr, to study genome wide patterns of differentiation and to look for candidate genes related to morph separation, we screened for genetic variation in developmental transcriptomes of three Þingvallavatn morphs (SB, LB and PL-charr) (Guðbrandsson et al. 2018). Because of the extra whole genome duplication in the ancestor of salmonids (Allendorf and Thorgaard 1984; Moghadam et al. 2011) and as each sample was a pool from three individuals, we developed a F-statistic based filter (*F*_*PT*_, see methods) to remove spurious variants caused by sequence divergence of paralogous genes. Variants with similar frequency in all samples most likely reflect sequence differences between stably expressed paralogs, but true polymorphisms should differ in frequency among samples. Simulations confirmed this assumption (see Figure S1), and from them we decided on *F*_*PT*_ = 0.1 as a threshold. It should be noted that paralogs differing in expression levels between morphs and/or timepoints escaped this *F*_*PT*_-filter and some variants in the dataset could be of that nature, thus a subset of variants was subject to validation (see below). After filtering, 19,252 variants remained, in 8,961 transcripts of 7,968 genes (Tables 1 and S1 and Supplementary file S2). As the data came from genetically related samples (each sample a pool of 3 embryos, from families produced by multiparent crosses), we could not apply standard population genetic analyses. Instead, we conducted principal component analyses, calculated *F*_*ST*_ between groups, tested for GO enrichment and mapped variants to linkage groups in order to characterize the patterns of genetic variation.

**Table 1:** The number of variants after each filtering step in the variant-calling pipeline.

|  | Variants |
| --- | --- |
| Freebayes first run | 143,744 |
| Cov and freq filters | 59,292 |
| $F_{PT}$ filter | 23,408 |
| Freebayes second run | 19,575 |
| Final set | 19,252 |

The three morphs separated at the genetic level. The first principal component (based on all variants) distinguished the planktivorous morph (PL-charr) from the two benthic morphs (LB- and SB-charr), which in turn separated on the second axis (Figure 2 A). The first two axes explain 19.5% and 12.7% of the variation, and other axes 4.2% at the most each (data not show). As the samples came from pooled crosses with 5-10 parents of each sex (see methods), the individuals sequenced may be more related than a random sample from the lake. But the different crosses for PL- and SB-charr cluster together in the first two PC-axis indicating that the separation of morphs is not an artifact of the experimental setup (see 2 A). To further investigate the genetic difference between morphs we studied variants with high *F*_*ST*_-values and private alleles in each morph (note that due to the sequencing of pooled samples from sibling embryos, the *F*_*ST*_ could be inflated). Using *F*_*ST*_ = 0.2 as a cutoff yielded 2331 variants (Figure 2 B). Variants were categorized by the magnitude of the absolute allele frequency difference between one morph and the other two, and called LB-, SB- or PL-charr specific variants. For instance, the SB-specific variants differed most strongly in allele frequency in SB-charr versus the average allele frequency of LB- and PL-charr. A significant excess (*χ*^2^=140.3, df = 2, p < 0.0001) of variants (1174) belonged to the PL-specific category (separating PL-charr and the two benthic morphs), while 605 and 552 variants associated with SB- and LB-charr, respectively.

**Figure 2:**
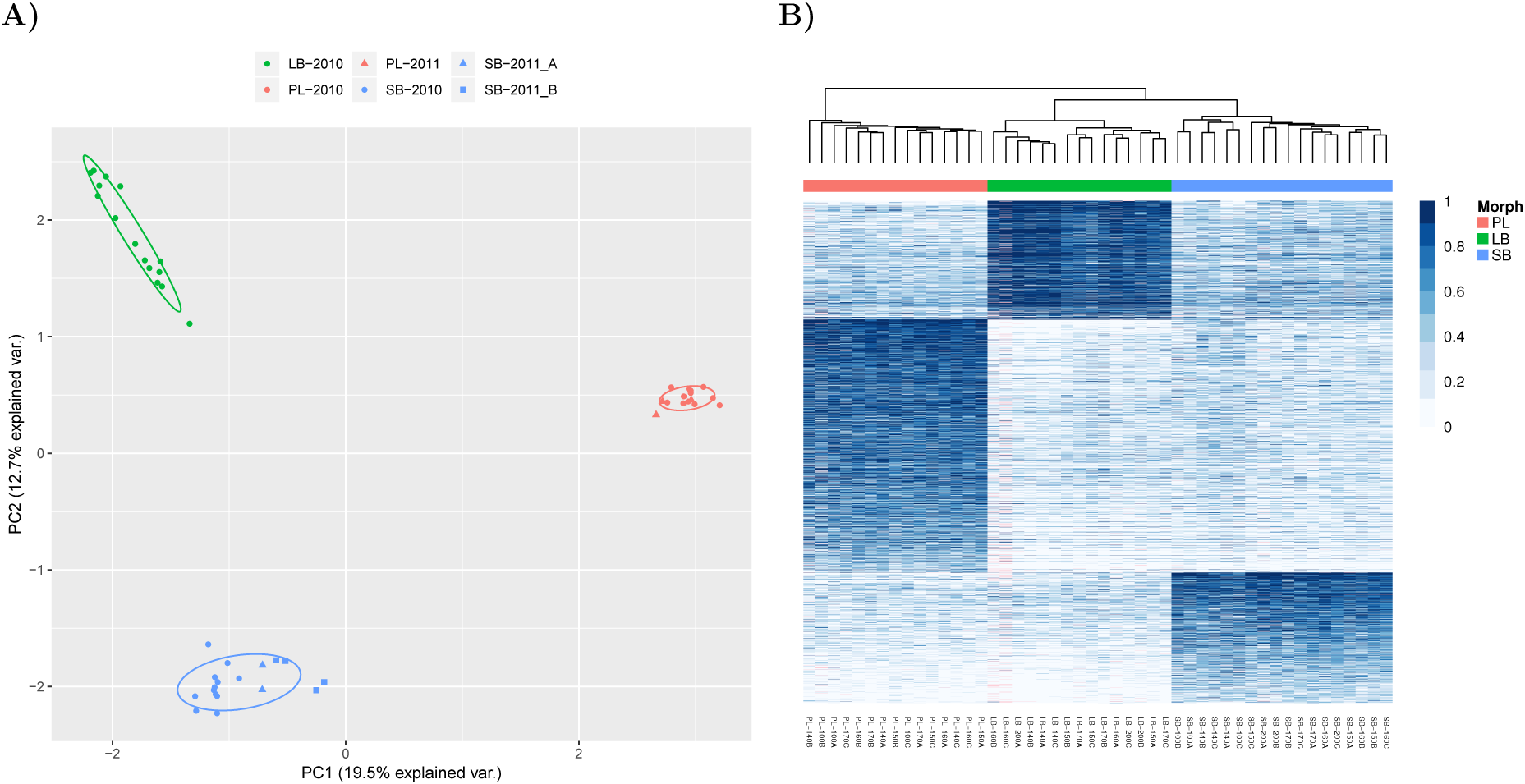
A) Genetic separation of samples from three sy1 mpatric Arctic charr morphs, based on principal component analysis of 19,252 transcriptome variants. The first and second principal components are depicted with the proportion of variance explained. Colours and shapes indicate which morphs (SB: blue, LB: green and PL: red) and crosses the embryos originated from (see legend). Note, while the samples from the three SB crosses do not overlap entirely, then they are very distinct from the PL and LB specimens. Overlaid are 68% normal data ellipses for each morph. **B)** Separation of morphs based on the 2331 variants with *F*_*ST*_ *>* 0.2. Each column represents a sample, name below indicates morph, developmental timepoint and biological replicate. The white-blue scale depicts allele frequency, higher frequency of the alternative allele with darker blue. Hierarchical clustering grouped samples and variants. This separated the morphs (abscissa) and similar variants by allele frequency (ordinate). A total of 1174 variants had a higher frequency in PL-charr, 552 in LB- and 605 in SB-charr. Missing values are indicated by pink. Coloring of individuals by morph, SB: blue, LB: green and PL: red.

A more stringent *F*_*ST*_ cut-off (0.4) exacerbated the PL-charr vs. benthic division (Figure S4). A higher fraction of variants (*χ*^2^=56.6, df=2, p > 0.0001) separated PL-charr from the other two (202 PL-specific, 71 SB-specific, and 51 LB-specific variants). The same pattern was observed for private variants (alleles with frequency <1% were considered absent from a morph), 56, 18 and 13 were private to PL-, SB- and LB-charr, respectively (Table S3). Note, differences in allele frequencies were between groups, and not evolutionarily polarized (ancestral versus derived). Based on these data we concluded that the largest genetic separation of the sympatric charr in Lake Þingvallavatn was between PL-charr and the two benthic morphs.

### 3.2 Genetic differences between benthic and limnetic morphs in collagen metabolism and environmental sensing

In order to gauge if certain biological systems differed between the morphs, we tested for GO-enrichment in variants associating with specific morphs. As some variants were annotated to different transcript isoforms of the same gene, testing was done both on transcripts and gene level using annotation of salmon genes. No GO categories were significant for SB-specific and LB-specific variants. However, PL-specific variants were significantly enriched in 10 GO-categories (Table 2). Nine categories were found at the transcripts level and seven on the gene level. Only one of the gene level categories significant was not at the transcript level (*collagen fibril organization*) but it was close to being significant (*FDR* = 0.015). Also, variants were enriched in one other category related to the extracellular matrix *collagen catabolic process*. That may have led to two higher level categories to be significant at the transcript level but not at the gene level (*multicellular organismal catabolic process* and *macro-molecule metabolic process*). Interestingly, four of the categories relate to environmental sensing and responses, *i*.*e*. light and sound, and showed a strong signal both on transcript and gene level, *e*.*g. inner ear morphogenesis* and *visual perception*. The final two categories were *tooth mineralization* and *odontogenesis*, both related to tooth development. GO-enrichment tests on private variants only revealed three GO-categories related to tRNA aminoacylation significant at the transcript level for variants private to PL-charr. The signal is most likely because three transcripts are annotated to the same gene rather than being a real biological pattern (Table S4). Together, the GO analyses of biological functions point to genetic differences in sensing, collagen metabolism and mineralization, between benthic and limnetic morphs. This could be linked to distinct differences in the main feeding habitats and principal prey species of PL-versus LB- and SB-charr and to differences in craniofacial features of these morphs.

**Table 2:** Gene ontology categories enriched in variants with *F*_*ST*_ above 0.2 and highest mean allele frequency deviation for PL-charr on the transcripts (*tr*) and gene (*ge*) level. The number of transcripts and genes with high divergence is shown (*PL*_*tr*_ and *PL*_*ge*_) and the total number of transcripts and genes tested in the category as well (*T ot*_*tr*_ and *Tot*_*ge*_). The multiple testing corrected p-value or false discovery rate (*FDR*) is also shown for both levels.

| Category | Term | $PL_{tr}$ | $Tot_{tr}$ | $FDR_{tr}$ | $PL_{ge}$ | $Tot_{ge}$ | $FDR_{ge}$ |
| --- | --- | --- | --- | --- | --- | --- | --- |
| GO:0007601 | visual perception | 30 | 122 | 0.0020 | 32 | 114 | 0.0014 |
| GO:0050953 | sensory perception of light stimulus | 30 | 122 | 0.0020 | 32 | 114 | 0.0014 |
| GO:0007605 | sensory perception of sound | 34 | 156 | 0.0025 | 36 | 137 | 0.0014 |
| GO:0034505 | tooth mineralization | 10 | 18 | 0.0025 | 9 | 13 | 0.0031 |
| GO:0030574 | collagen catabolic process | 23 | 89 | 0.0069 | 20 | 60 | 0.0071 |
| GO:0042472 | inner ear morphogenesis | 30 | 137 | 0.0083 | 32 | 128 | 0.0087 |
| GO:0030199 | collagen fibril organization | 19 | 73 | 0.0150 | 18 | 51 | 0.0073 |
| GO:0044243 | multicellular organismal catabolic process | 23 | 93 | 0.0087 | 20 | 65 | 0.0140 |
| GO:0042476 | odontogenesis | 26 | 109 | 0.0083 | 24 | 92 | 0.0359 |
| GO:0043170 | macromolecule metabolic process | 499 | 4,790 | 0.0083 | 490 | 4,155 | 0.3650 |

### 3.3 Genome-wide divergence of sympatric charr morphs

To explore if the genetic differences between morphs were restricted to specific chromosomal regions or more widely distributed, we mapped the variants to the Arctic charr genome using `blastn` (see methods). A large fraction (93%, 17,933 of 19,252) mapped to the genome. Of those variants 10,956 or 61% mapped to chromosomes but the rest to unplaced scaffolds and contigs. The number of variants mapped on each linkage group was, as expected, related to the linkage group size (Figure S2). Markers with *F*_*ST*_ > 0.2 were found on all linkage groups, and peaks of more pronounced differentiation were found on many chromosomes. Similarly, the three types of high *F*_*ST*_ markers (PL-, SB- and LB-charr specific) were found on almost all linkage groups (Figure 3 B).

**Figure 3:**
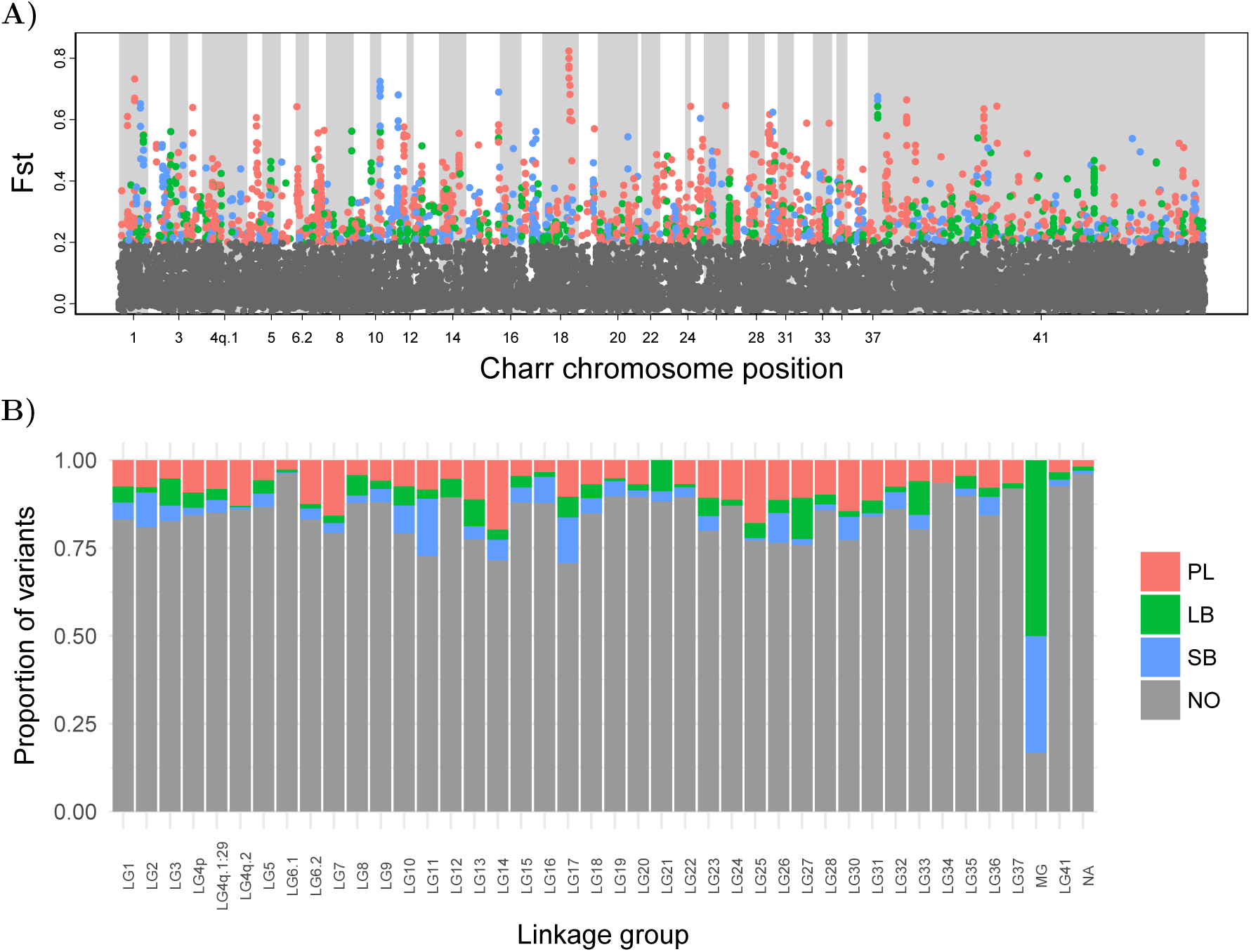
A) *F*_*ST*_ values plotted by position of variants on the Arctic charr genome from (Christensen, Rondeau, Minkley, Leong, Nugent, Danzmann, Ferguson, Stadnik, Devlin, Muzzerall, Edwards, Davidson and Koop 2018). The colors indicate which morph differs most strongly in allele frequency from the other two for variants with *F*_*ST*_ above 0.2 (SB: blue, LB: green and PL: red) and gray represents variants with *F*_*ST*_ below 0.2. **B)** Proportion of each variant group on each linkage group (MG: mitochondrial chromosome). Unplaced scaffolds and contigs are represented by “linkage” group 41 and in **B)** NA refers to unmapped markers.

There were indications of enrichment of high *F*_*ST*_ variants of a particular type in particular linkage groups, most dramatically on the mitochondrial chromosome. Previously we found genetic differentiation between SB-charr and Aquaculture charr in the mtDNA (Gudbrandsson et al. 2016), using transcriptome data and genotyping of population samples. There we observed a high frequency of derived alleles in LB- and SB-charr. In the current dataset, 6 variants were observed in the mtDNA (Table S1) and all but one had high *F*_*ST*_ (Figure 3). Consistent with previous results, these variants were LB- or SB-specific and three of those (m1829G>A, m3211T>C and m3411C>T) were within the 12s and 16s rRNA genes (Gudbrandsson et al. 2016). A lower fraction of markers had high *F*_*ST*_’s on other linkage groups. Distinct chromosomal regions harboured variants with high differentiation associating with particular morphs. For instance, a high peak of PL-specific variants was observed on LG18, and a peak of SB-specific markers on LG10 (Figure 3) and on LG1 high *F*_*ST*_ variants of all categories (PL, SB, LB-specific) were found. In sum, the genetic separation between morphs was found on all linkage groups, including strong differences in the mitochondrial DNA.

### 3.4 Verification of variants in population sample confirms morph separation and regions of differentiation

As the candidate variants were derived from sequenced pools of embryos from multi-parent families, we wanted to verify them in population samples from the wild to address three questions: First, do estimates of allele frequencies in the transcriptome and in wild populations correspond? Second, do the three morphs differ genetically in population samples? Third, do some markers associate fully with morph status? Fourth, how broad are the regions of differentiation? Fifth, does the understudied Piscivorous (PI) charr differ genetically from the other three morphs?

Candidate variants (23 in total, Table S5) were chosen based on high *F*_*ST*_ values in the transcriptome sequencing, chromosomal location, biological functions and/or differential expression (Guðbrandsson et al. 2018). Sexually mature individuals of all four morphs in Lake Þingvallavatn were genotyped. All markers but one amplified successfully and behaved as true single nucleotide polymorphisms (SNPs, Table S5). The allele frequencies and estimated *F*_*ST*_ values in the transcriptome and population sample were highly correlated (Kendall’s *τ* = 0.76, *p <* 0.0001, and *τ* = 0.58, *p <* 0.001 respectively) (Figure S5). Allele frequencies seemed to be underestimated in the transcriptome, particularly lower frequencies (< 0.3) (Figure S5 A). The overall *F*_*ST*_’s for individual markers ranged from 0.01 (*gnl3l_G795T*) to 0.66 (*gas1l_A3641C* and *wee1_T305A*) (Table S7). The correspondence of allele frequencies in the transcriptome and the population sample, suggests the large scale patterns in the former are real. Majority of markers (18/22) deviated significantly from Hardy-Weinberg equilibrium when tested on the 93 samples (all four morphs), but almost all were in HWE within each morph (Table S7). Consistently, the majority of the variance in genotypes associated with morph (Figure 4 A). PC1 (45.2% of variance explained) separated benthic-limnetic morphs while PC2 (14.4%) distinguished three morphs (SB, LB and PL-charr). Note, the higher proportion of variance explained by the first two PC’s for the genotyped SNPs (compared to transcriptome) likely reflects the non-random selection of variants with large frequency differences for genotyping.

**Figure 4:**
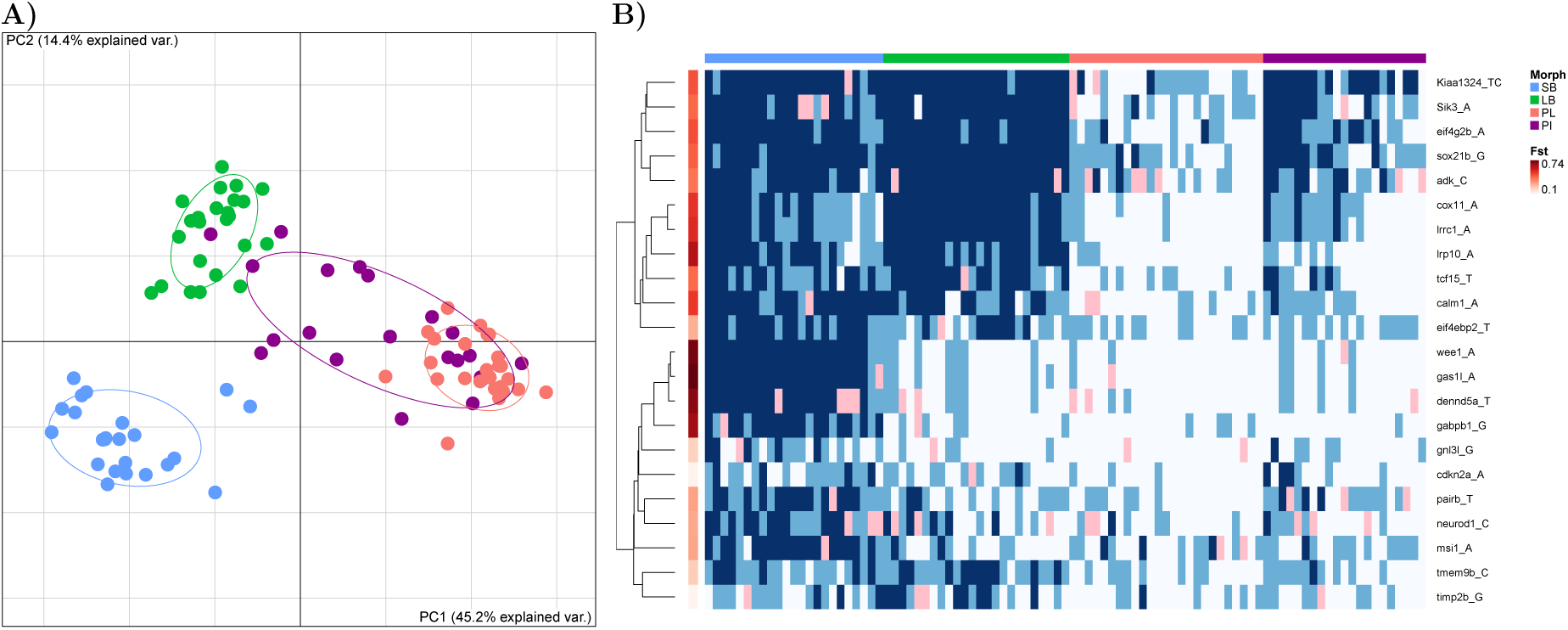
A) Genetic separation of the four morphs sympatric charr morphs depicted with principal component analyses of 22 variants genotyped in population samples. Individuals are graphed according to scores of the first two PC’s, along with 68% normal data ellipses for each morph (SB: blue, LB: green, PL: red and PI: purple). **B**) Heatmap of genetic variation in population samples of four morphs, depicting association of variants with morphology and linkage of markers. Genotypes of the 22 variants were clustered by genes. *F*_*ST*_ values (shades of red) for morphs are graphed for each marker. The morphs are color coded as in **A)**, and genotypes, homozygous reference allele (white), heterozygous (light-blue) and homozygous alternate allele (blue), with pink indicating missing data.

Eleven markers that separated benthic-limnetic morphs were genotyped (Figure 4 B), but despite strong associations, no marker coupled fully to a morph or morphotype. The 22 genotyped variants mapped to 12 separate linkage groups and 3 unplaced scaffolds (Supplemental table S5). Note that the resolution of the data is limited. For instance, the region around *calm1* on LG4.q2 had numerous variants with high differentiation between PL and the benthic morphs in the transcriptome (Figure 5 A), but it is unclear whether that reflects a single or many differentiated regions.

**Figure 5:**
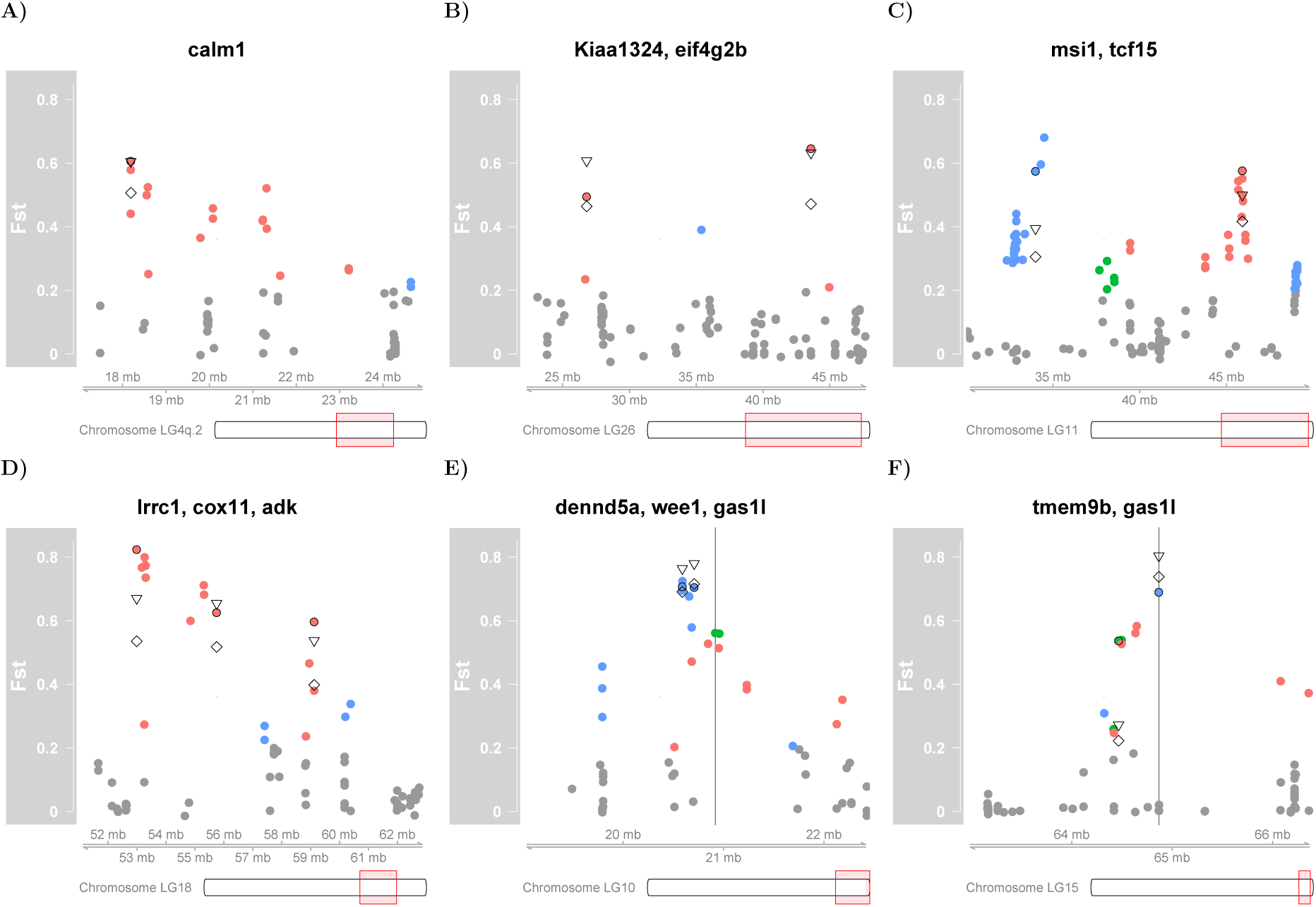
Detailed view of chromosomal regions and variants showing high divergence in the KASP-assay. **A)** Region on LG4q.2 that includes *calm1* and many other variants with high *F*_*ST*_ values (not validated). **B)** A large region on LG26 that includes *Kiaa1324* and *eif4g2b*. **C)** A region on LG11 that includes *msi1* and *tcf15*. **D)** A region on LG18 that includes *lrcc1, cox11* and *adk*. **E)** A region on LG10 that includes *dennd5a, wee1*. The location of the second best blast hit for the variant in *gas1* is shown with a vertical line. **F)** A region on LG15 that includes *tmem9b* and *gas1l*. The location of the best blast hit for the variant in *gas1* is shown with a vertical line. The regions on LG10 and LG15 shown in **E)** and **F)** are ohnologous and some of the markers might belong in the opposite linkage group as we suspect is the case for *gas1l*. Colored dots are values from the transcriptome as in Fig. 3 and validated variants are marked with a black circle (∘). The triangles (∇) show the *F*_*ST*_ value from the KASP-assay for three morphs (PL, SB and LB) and the diamonds (⋄) *F*_*ST*_ for all morphs (including PI).

To ask whether there were multiple independent regions of differentiation within the same chromosome we chose several supposedly linked variants (from four specific linkage groups) from the transcriptome for genotyping in the population samples and for linkage disequilibrium (LD) within each morph. Note, the LD-estimates must be interpreted cautiously as they are imprecise because only 21 or 24 fish were sampled from each morph (Yan et al. 2009).

We surveyed four sets of putatively linked markers. Two pairs of markers were separated by more than 10 Mb, and showed very low or no association. These were PL-specific markers (*Kiaa1324_TC393AA* and *eif4g2b_G652A*) on LG26 (Figure 5 B) and two markers in (*msi1* and *tcf15*) on LG11 (Figure 5 C, Table 3).

**Table 3:** Estimates of LD (*r*^2^), within each morph, and genomic distance for pairs of genotyped variants that mapped to the same linkage group or showed a strong association.

| Var1 | Var2 | Pattern of divergence<sup>a</sup> | Linkage group | Genomic distance (bp) | SB | LB | PL | PI |
| --- | --- | --- | --- | --- | --- | --- | --- | --- |
| *lrrc1* | *cox11* | PL | LG18 | 2,762,681 | 0.90 | 1.00 | 0.73 | 0.90 |
| *cox11* | *adk* | PL | LG18 | 3,370,151 | 0.04 | NA | 0.06 | 0.23 |
| *lrrc1* | *adk* | PL | LG18 | 6,132,832 | 0.03 | NA | 0.00 | 0.32 |
| *eif4g2b* | *Kiaa1324* | PL | LG26 | 16,793,974 | 0.08 | 0.00 | 0.00 | 0.06 |
| *msi1* | *tcf15* | SB, PL | LG11 | 11,914,739 | 0.02 | 0.06 | 0.01 | 0.00 |
| *dennd5a* | *wee1* | SB | LG10 | 118,070 | 0.82 | 0.92 | 0.52 | 1.00 |
| *wee1* | *gas1l* | SB | LG10<sup>b</sup> | <sup>b</sup>210,891 | 0.51 | 0.85 | 0.51 | 0.95 |
| *dennd5a* | *gas1l* | SB | LG10<sup>b</sup> | <sup>b</sup>328,961 | 0.42 | 0.79 | 0.47 | 0.95 |
| *gas1l* | *tmem9b* | SB, PL | LG15 | 400,866 | 0.02 | 0.00 | 0.00 | 0.26 |

<sup>a</sup> In which morph did the variant differ most in frequency, first description for Var1 and second for Var2.
<sup>b</sup> Second best blast hit for *gas1l* was on LG10, the best was on LG15.
NA: Not available, because variant *adk* was not polymorphic in LB.

More tightly spaced were three markers mapping to LG1 that separated PL-charr from the benthic morphs (Figure 5 D). While *lrrc1* and *cox11* are 2.8 Mb apart the markers showed tight LD (*r*^2^ > 0.72) (Table 3) but the third marker (*adk_C1075T*), located 3.4 Mb downstream of *cox11*, had weak or no LD with the other two (Table 3). This suggests there are two differentiated regions within this ∼6 Mb region of the genome.

Three SB-specific variants (*gas1l_A3641C, wee1_T305A* and *dennd5a_A2555T*) on LG10 were in strong LD (*r*^2^ *>* 0.45) in all morphs. While the variants in *dennd5a* and *wee1* mapped 118 kb apart, then the region containing the *gas1l* variant aligned to LG15 (The second best hit was on LG10 close to the other two, Figure 5 E). LG10 and LG15 are ohnologous and have over 90% sequence similarity (Christensen, Rondeau, Minkley, Leong, Nugent, Danzmann, Ferguson, Stadnik, Devlin, Muzzerall, Edwards, Davidson and Koop 2018). Notably, the *gas1l* variant was in linkage equilibrium with another variant on LG15 (in *tmem9b*, Figure 5 F, Table 3). These discrepancies may be due to assembly errors in the Arctic charr draft genome by Christensen, Rondeau, Minkley, Leong, Nugent, Danzmann, Ferguson, Stadnik, Devlin, Muzzerall, Edwards, Davidson and Koop (2018) as the regions with the three SB-specific variants all mapped to the same chromosome in *S. salar* (ssa26) and *O. mykiss* (omy06) genomes. Otherwise, there was low LD between variants (Table S8).

Finally, the phyletic relationship of the rare PI-charr to the other morphs is unknown (Gíslason 1998; Volpe and Ferguson 1996). It was hypothesized that PI-charr could be an ontogenetic morph of PL-charr that learn to utilize threespined sticklebacks as prey (Snorrason et al. 1989). No transcriptome data were available for PI-charr, but we genotyped 21 individuals. Contrary to the other three morphs, PI individuals did not group in the PC plot (Figure 4 A). This may depend on some technical shortcomings, i) too few markers, ii) ascertainment bias or iii) incorrect classification of PI-individuals. If technical biases do not dominate, then the data may reflect biological patterns. At face value, the data suggest the PI-charr are more heterogeneous genetically than the other morphs. And the data do not support the ontogenetic shift hypothesis (which predicted the PI should cluster with the PL-charr). Curiously, half of the PI-charr were in the PL-cluster but the rest (except one) fell in between the PL- and LB-clusters. Thus while the other three morphs separated clearly genetically, determination of the genetic status and nature of the piscivorous charr requires more data.

In summary, while no single variant associated fully with specific morph many distinct regions within chromosomes harbour variants separating the morphs.

## 4 Discussion

The four sympatric morphs of Arctic charr in Lake Þingvallavatn differ in a host of phenotypic attributes, e.g. in adult morphology, habitat choice, feeding preferences, growth pattern, size and age at maturity (see references in Sandlund et al. 1992). Common garden rearing experiments indicate these differences are based on genotype but also influenced by the environment, and emerge during embryonic and juvenile development (Skúlason, Noakes and Snorrason 1989; Parsons et al. 2010; Kapralova et al. 2015). While population genetics indicated a genetic separation of PL-, LB- and SB-charr (Magnusson and Ferguson 1987; Volpe and Ferguson 1996), the degrees of relatedness and phyletic relationships of these morphs have been unresolved. Here we utilized SNPs from RNAseq data derived from embryos of pure crosses of three morphs (Guðbrandsson et al. 2018) to assess their genetic separation. For each variant, we asked if one of the morphs was more diverged, and if so, how morph-specific variants were distributed in the genome and whether they related to particular functional systems. Although the experimental design, pools of individuals from bulk crosses of each morph, was not optimal for these purposes the data yielded many informative variants. Consistently, 22 of the 23 selected variants genotyped in independent population samples were real variants. Estimates of *F*_*ST*_ in the transcriptome may have been inflated relative to the population samples (15 out of 22 markers had lower *F*_*ST*_ in the latter, Figure S5). Notably, the frequencies of rare alleles were underestimated in the transcriptome as expected (Konczal et al. 2014) but might be exaggerated due to the sequencing of related individuals, pooling of three embryos in individual samples or Beavis effects (Beavis 1994) because high *F*_*ST*_ SNPs were chosen for verification. These facts limit the interpretation of this dataset, precluding for example estimation of the site frequency spectrum. But they do not negate the observed widespread genetic differentiation between morphs, as the correspondence of allele frequencies estimated with the two methods used (RNAseq and KASP-assay) and the genetic differentiation of morphs is stronger than the variation between crosses within the same morph, that suggest the patterns are biological and not a side effect of the experimental design.

Previously, we (Gudbrandsson et al. 2016) extracted variants from RNAseq data from SB-charr of Þingvallavatn and an aquaculture breeding stock derived from several populations (mainly from the north of Iceland (Svavarsson 2007) - an outgroup for Þingvallavatn charr). While that study also yielded ∼20,000 candidate variants, the proportion of variants with high *F*_*ST*_ was larger, probably reflecting more divergence between the aquaculture charr and the SB-charr than among the three Þingvallavatn morphs studied here. It is unknown how many of the variants observed in these RNAseq data are shared with other populations in Iceland and across the species range (Brunner et al. 2001).

### 4.1 Genetic separation of recently evolved sympatric morphs

Geological forces, volcanic activity, isostatic rebound as well as underground rivers and springs in the rift-zone generate specific niches for the Icelandic biota (Kornobis et al. 2010; Kristjánsson et al. 2012; Marteinsson et al. 2013; Gudmundsdóttir et al. 2017), but also perturb them. Lake Þingvallavatn formed ∼10,000 years ago but resides in a geologically active rift zone, with the most recent eruption ∼2000 years ago (Saemundsson 1992). Furthermore, the tempo of the Ice-age glacial retreat was uneven, interleaved with periods of glacial advance (Saemundsson 1992; Norðdahl et al. 2008). Thus, the history of colonization and adaptations by Arctic charr and other organisms in this geographic region may be quite complex.

Studying the relatedness of the morphs can shed some light on the history of colonization and divergence in Lake Þingvallavatn. The morphs (PL-, LB- and SB-charr) are more closely related to one another than to other Icelandic populations (Kapralova et al. 2011). The data presented here firmly reject hypothesis I, that the PL-, LB- and SB-charr morphs are formed by environmental influence on individuals of a shared gene pool (the PI-charr may be an exception, see below). The data also suggest that the benthic morphs (LB- and SB-charr) are more closely related. This is congruent with patterns of differential expression in this transcriptome (Guðbrandsson et al. 2018) and one previous study (Volpe and Ferguson 1996) but not others (Magnusson and Ferguson 1987; Danzmann et al. 1991; Gíslason 1998). This incongruence and the observed distribution of the high *F*_*ST*_ variants into PL-, LB- and SB-specific categories may reflect limited resolution due to few markers in earlier studies of Lake Þingvallavatn charr, incomplete sorting of alleles or differential effects of selection on distinct traits and variants. Coalescence simulations based on microsatellite data were more consistent with a brief phase of allopatric separation of the ancestors of PL- and SB-charr Lake Þingvallavatn, rather than sympatric evolution (Kapralova et al. 2011). This study considered two demographic scenarios and two parameters (migration and population size). Analyses of other scenarios, *i*.*e*. with changing migration rates or introgression would be most interesting (see for example Jacobs et al. 2018). Considering the episodic nature of ice retreat and the high geological activity in the Þingvallavatn area during the formation of the lake (Saemundsson 1992; Norðdahl et al. 2008) charr may have invaded the waters more than once, or experienced intromittent isolation of populations in rifts. The current study lacks a natural outgroup, and future population genomic surveys of these morphs should include populations from the neighbouring geographic area, including the Þingvallavatn outflow river Sog and its downstream Lake Úlfljótsvatn. Lake Úlfljótsvatn is particularly interesting as it also hosts landlocked morphs of charr, including forms similar to the PL- and SB-charr (Jóhannsson et al. 1994; Woods et al. 2012a).

The *Salvelinus* genus is particularly interesting for sympatric polymorphism (Noakes 2008; Klemetsen 2010; Muir et al. 2016; Markevich et al. 2018) that seems to run counter to the predominant mode of allopatric speciation. Smith and Skúlason (1996) argued resource polymorphism and phenotypic plasticity could potentiate ecological speciation, even within a geographic area. The question remains if charr morphs in sympatry (e.g. in the Transbaikalian lakes (Gordeeva et al. 2015), Lake Galtarból in northern Iceland (Wilson et al. 2004), and Lower Tazimina Lake in Alaska (May-Mcnally et al. 2015)) differentiated in true sympatry, or originate by repeated invasions of the water bodies. The morphs in Loch Stack (Adams et al. 2008) and Loch Tay (Garduño-Paz et al. 2012) are clearly of allopatric origin. A recent genome-wide study of sympatric charr in Scotland and Siberia is consistent with sympatric origins of charr morphs (Jacobs et al. 2018). Ecological specialization can lead to differential growth, behavior and maturity that may influence both choice of spawning sites and timing of spawning. Within Lake Þingvallavatn, the timing of spawning differs between the three best-studied morphs. The LB-charr spawns in August, PL-charr in October, but SB-charr over a broader period from September to November (PI-charr spawn in October) (Skúlason, Snorrason, Noakes, Ferguson and Malmquist 1989; Sandlund et al. 1992). Although all morphs appear to spawn in the same habitat, i.e. among loose stones in the littoral zone, there may be micro-habitat preferences for spawning sites. Variation in mating behavior can in principle lead parents to choose specific nesting sites, e.g. close to inflow of cold water. Such differences in environmental conditions could, in turn, influence development of ecological functional traits and adult form of the morphs.

The ambiguous genetic status of the large, rare and piscivorous morph is quite intriguing. PI-charr was only studied by genotyping 22 markers, but curiously PI individuals did not cluster as tightly as the other morphs. PI individuals grouped with LB, PL-charr and a few in between. Previous studies found PI individuals affiliated with either or both morphs (Magnusson and Ferguson 1987; Volpe and Ferguson 1996). Several scenarios may explain the data and the origin of PI-charr. First, classification of the morph based on phenotype may not be stringent enough and thus some LB-charr might have been misidentified as PI-charr. As the LB- and PI-charr utilize different prey and differ in spawning periods (the former in August and later in October (Skúlason, Snorrason, Noakes, Ferguson and Malmquist 1989)) this misidentification is unlikely. However, it is possible that some LB-charr males might be erroneously classified as PI because of an extended jaw-hook, as some are still running in October. Secondly, it is possible that PI-charr is genetically distinct, but that the markers chosen for genotyping may not suffice to distinguish them. Thirdly, Snorrason et al. (1989) postulated that PI-charr emerges as certain PL-charr learn to eat fish. PI-charr would then be environmentally induced, genetically identical to PL-charr. The fact that PI-charr does not overlap exclusively with PL-charr argues against this theory. Fourthly, PI-charr may actually be more heterogeneous at the genetic level than the other three morphs. This might reflect recent origin or perhaps a combination of ontogenetic shift of some individuals from PL stock with repeated hybridizations with other morphs (most likely LB-charr because of the size). Currently, we can not distinguish between these possibilities.

We conclude that three of the Lake Þingvallavatn morphs differ genetically, and considering the differences in trophic traits and spawning, may even be reproductively isolated (hypothesis III). It remains to be determined if the PI-charr are genetically separated from the other morphs or if they are an environmentally induced form as may be the case of eco-morphs in Sockeye salmon *Oncorhynchus nerka* (Limborg et al. 2018).

### 4.2 Genome-wide separation of sympatric morphs

How extensive is the genetic separation of morphs and how is the genetic differentiation distributed in the genome? For instance, are differentiating variants localized to a few islands (Nadeau et al. 2012; Andrew and Rieseberg 2013; Malinsky et al. 2015) as stated in hypothesis II or are there multiple signals on many chromosomes (Hohenlohe et al. 2010; Jones et al. 2012) according to hypothesis III?

Variants with relatively high *F*_*ST*_ (>0.2) mapped to all 40 *S. alpinus* linkage groups (including the mitochondrial chromosome), which supports hypothesis III. The clustering of variants, some with *F*_*ST*_ >0.5, suggests specific genes/regions associated with the specializations of particular morphs. Analyses of LD between genotyped variants implied that peaks on the same linkage group were independent. The distribution of variants on linkage groups was rather even, except the mtDNA had a high fraction of LB- and SB-specific variants (consistent with earlier findings (Gudbrandsson et al. 2016)). This might reflect history or adaptive evolution of mitochondrial functions in the benthic morphs. The extensive genetic separation of these three morphs suggests many differentiating genes on many (or all) linkage groups. While the data seem to refute hypothesis II, that few genomic islands differentiate the morphs, it is plausible that ecological specialization has driven differentiation in certain genomic regions.

Previous studies of salmonids have found genome-wide differences between populations and ecological morphs. Significant genetic differences were found between migratory and non-migratory Rainbow trout (Hale et al. 2013), the lake, river and stream ecotypes of sockeye salmon (Larson et al. 2017) and ecologically different subpopulations of salmon (Vincent et al. 2013; Cauwelier et al. 2017). Sympatric ecotypes are found in several salmonid species, e.g. whitefish (Gagnaire et al. 2013), but are most pronounced in Arctic charr and Lake charr *Salvelinus namaycush*. Genomic studies of the latter revealed varying degrees of genetic separation of sympatric morphs in large lakes in North America (Harris et al. 2015; Perreault-Payette et al. 2017). A study on five pairs of benthic-limnetic whitefish morphs (Gagnaire et al. 2013) found a correlation between genetic and phenotypic differentiation and considerable overlap of genomic regions that differentiated morph pairs in each lake. Notably, the significant degree of genetic parallelism broke up at the finer level, with different haplotypes associating with morphotype, suggesting genetic and/or allelic heterogeneity of the causative loci (Gagnaire et al. 2013). Recently, genome wide analyses of Scottish and Siberian Arctic charr suggested limited genetic parallelism in benthic-limnetic specializations (Jacobs et al. 2018). Fine scale analyses of peaks of differentiation could not be conducted with the current data, but the genome wide distribution of differentiated variants suggests restricted gene flow between at least three of the morphs (PL-, LB- and SB-charr) and that they may be reproductively isolated.

Genomic differentiation reflects the history and evolution of groups, degree and age of separation of groups, extent of gene flow and hybridization events (Seehausen et al. 2014; Shapiro et al. 2016). But intrinsic factors, such as genes and their biological actions, chromosomal inversions and polymorphisms, also shape the rates of differentiation and divergence, and genomic features, such as centromeres, gene density, recombination and GC content influence estimates of sequence divergence and allele frequency differences (Seehausen et al. 2014; Burri et al. 2015; Sutherland et al. 2016; Vijay et al. 2017). Population genomic data and analyses are needed to disentangle the role of positive selection and intrinsic factors on patterns of differentiation in this system. More broadly, the confirmed synteny of large genomic regions (Nugent et al. 2017), the range of species, subspecies and ecologically distinct populations (Klemetsen 2013; Jacobs et al. 2018) sets the stage for future studies of the genomic and ecological correlates of divergence and polymorphism in salmonids.

### 4.3 Potential mechanisms of phenotypic and developmental differences between sympatric morphs

Which genes influence the many phenotypic differences between the morphs? Genetic variation in transcribed sequences tags large fraction of the potentially functional regions of the genome. The genome-wide differentiation between morphs implies polygenic basis of their differences, but major genes influencing specific traits may be segregating. The genomic resolution of the current data is low, and while we can describe frequency differences in variants in specific genes, it is more likely that other linked polymorphisms/loci are actually contributing to phenotypic differences.

The current data can inform future studies by bringing attention to particular chromosomal regions, genes and systems that may influence differences between charr morphs. The GO results suggest the limnetic and benthic morphs differ genetically in three systems (collagen metabolism, tooth mineralization and sensory functions). The enrichment of variants in collagen organization/catabolism and extracellular matrix categories is consistent with observed differential expression of ECM related and cartilage remodeling genes in benthic and limnetic morphs (Ahi et al. 2014, 2015). Strong differentiation was found for a variant in *calm1*. Calmodulins are broadly expressed in vertebrate tissues, including bone and articular cartilage in humans (Mototani et al. 2005), and notably higher expression of *Calm1* was found in beak primordia of finches with longer beaks (Abzhanov et al. 2006). Secondly, we found frequency difference of a variant in *eif4ebp1* between morphs (*F*_*ST*_ = 0.28). This gene had higher expression in SB compared to PL-charr in developing embryos (this transcriptome (Guðbrandsson et al. 2018)). Curiously, the same was seen in adult muscles of SB-charr from other locations in Iceland (Macqueen et al. 2011). *eif4ebp1* is involved in the insulin receptor signaling pathway and mTOR signalling (Bidinosti et al. 2010; Gkogkas et al. 2012; Banko et al. 2007). However, the *eif4ebp1* variant associated with benthic morphotype not small benthic charr, which is at odds with the proposition that reduced growth of the SB populations is mediated through changes in the mTOR activity. Finally, three SB-specific variants on linkage group 10 (or 15) had high *F*_*ST*_’s. This might represent long haplotypes with strong differentiation between morphs, that associate with the small benthic phenotype. Curiously one gene in the region (*Wee1*) is a key regulator of the timing of mitosis and cell size, first identified in fission yeast (Nurse and Thuriaux 1980) and another (*gas1, Growth arrest-specific protein 1*) associates with embryonic cranial skeleton morphogenesis and palate development (Seppala et al. 2007).

Few traits have been mapped to a gene in salmonids. The most celebrated exception being *vgll3* which associates with the age of maturity in wild populations, with very curious sex-dependent dominance of alleles (Barson et al. 2015; Ayllon et al. 2015). Unfortunately the *vgll3* gene was not transcribed in our data, but SB-specific variants with *F*_*ST*_ around 0.4 are found ∼1 Mb from the location of *vgll3* on LG2 (Figure S6 K). Further work is needed to check if variants in *vgll3, tulp4* (see Larson et al. 2017), *Greb1L* (see Prince et al. 2017) or other genes contribute to phenotypic differences in charr. QTL and fine-scale association mapping (Zimmerman et al. 2000; Palsson and Gibson 2004; Dworkin et al. 2005) are needed to find the causative variants. In sum, the data presented here can lead to the identification of genes influencing specific phenotypes separating the sympatric morphs, but more crucially suggest a polygenic basis for the morph differences.

### 4.4 Conclusion and future perspective

We mined genetic variation from three of the four morphs in Lake Þingvallavatn to address questions about their genetic separation, genome-wide differentiation of morphs and potential functionality of loci separating them. We started with three hypotheses about the putative causes of morph differences: Hypothesis I stated that the morphs are environmentally induced. According to hypothesis II, ecological specialization led to few genomic islands of strong genetic differentiation between morphs, with low background separation across the genome. Hypotheses III postulated substantial differentiation across the genome due to reduced gene flow, with the bulk of the genome showing separation between morphs. Estimates of genetic differentiation between the Þingvallavatn morphs have yield different results from low (Volpe and Ferguson 1996; Kapralova et al. 2011) to extremely high (Kapralova et al. 2013) depending on the marker used. Based on the genetic differentiation observed in this study, we conclude that three of the morphs (SB, LB and PL-charr) are distinct populations and find it highly unlikely that these morphs are environmentally induced (rejecting hypothesis I). Furthermore, we interpret the data as suggesting genome-wide differentiation over weak differentiation with a few islands of differentiation (hypothesis III over II). This is supported by the facts that the background *F*_*ST*_ in the transcriptome was rather high and high *F*_*ST*_ peaks were found on all linkage groups (and even multiple peaks on some). The fact that spawning times of certain morphs do not overlap (LB-charr spawn in August, but PL- and SB-charr in September - October) also argues that gene flow has been reduced between these morphs. Note however that hybrids of specific morphs, SB-, PL- and PI-charr can be generated in the laboratory (Kapralova 2014, Kalina H. Kapralova, Sigurdur S. Snorrason, Zophonías O. Jónsson, Arnar Pálsson *et al*. unpublished data) but their fitness and how common they are in nature is unknown. Thus we conclude that the three studied morphs in Lake Þingvallavatn are not one panmictic population and that gene flow between them has been limited. The nature of the PI-charr is still in doubt, some PI-charr may arise every generation by ontogenetic shift as suggested by Snorrason et al. (1989) while other PI may propagate, and even hybridize with the LB-charr. Population ddRAD-seq data of sexually mature/spawning charr of all morphs confirm the clear demarcation of three of the morphs and offer insights into the nature of the PI-charr (Han Xiao, Benjamín Sigurgeirsson, et al. unpublished data). Future studies of the fertilization success of hybrids and pure morphs, analyses of the development and fitness of hybrids and pure morph, and behavioural studies of spawning behaviour, reed locations and properties, and mate choice have can cast light on potential pre- and postzygotic barriers to gene flow between morphs. In the future, the phenotypic and genetic differences between sympatric and locally adapted Arctic charr populations can aid studies of ecological adaptation and the synthesis of evolutionary, ecological and developmental biology.

## 5 Acknowledgments

Special thanks to present and past members of the Arctic charr group at the University of Iceland for support, comments and suggestions. Thanks to Bjarni K. Kristjansson, Camille LeBlanc and colleagues at Holar University College for support. Thanks to Ben Sutherland and Eric Normandeau for excellent advice and open communication about MapComp and methods for placing variants on linkage maps. This project was supported by The Icelandic Center for Research (RANNIS #100204011) to SSS and coworkers, The University of Iceland Doctoral Fund to JG and University of Iceland research fund to AP, SSS and ZOJ.

## 6 Author contributions

Conceived and designed the study: JG, AP, ZOJ, SSS, SRF, KHK

Sampling: SSS, KHK, SRF, ZOJ, AP, JG

Gathered the data: ZOJ, SRF, KHK, VH, ÞMB

Retrieval and analyses of variants data: JG

SNP confirmation: KHK, AP, VH, ÞMB, JG

Analyses: JG, AP, KHK

Writing: AP, JG, SSS, KHK, SRF, ZOJ

## 7 Data access

The sequencing reads from the 48 RNA-seq samples were deposited into the NCBI SRA archive under BioProject identifier PRJNA391695 and with accession numbers: SRS2316381 to SRS2316428. Supplementary tables and files are available on figshare https://doi.org/10.6084/m9.figshare.c.4565735.v1 (specific hyperlinks in Supplementary material).

## Supplementary Material

### Supplementary Tables and Files

**Table S1:**
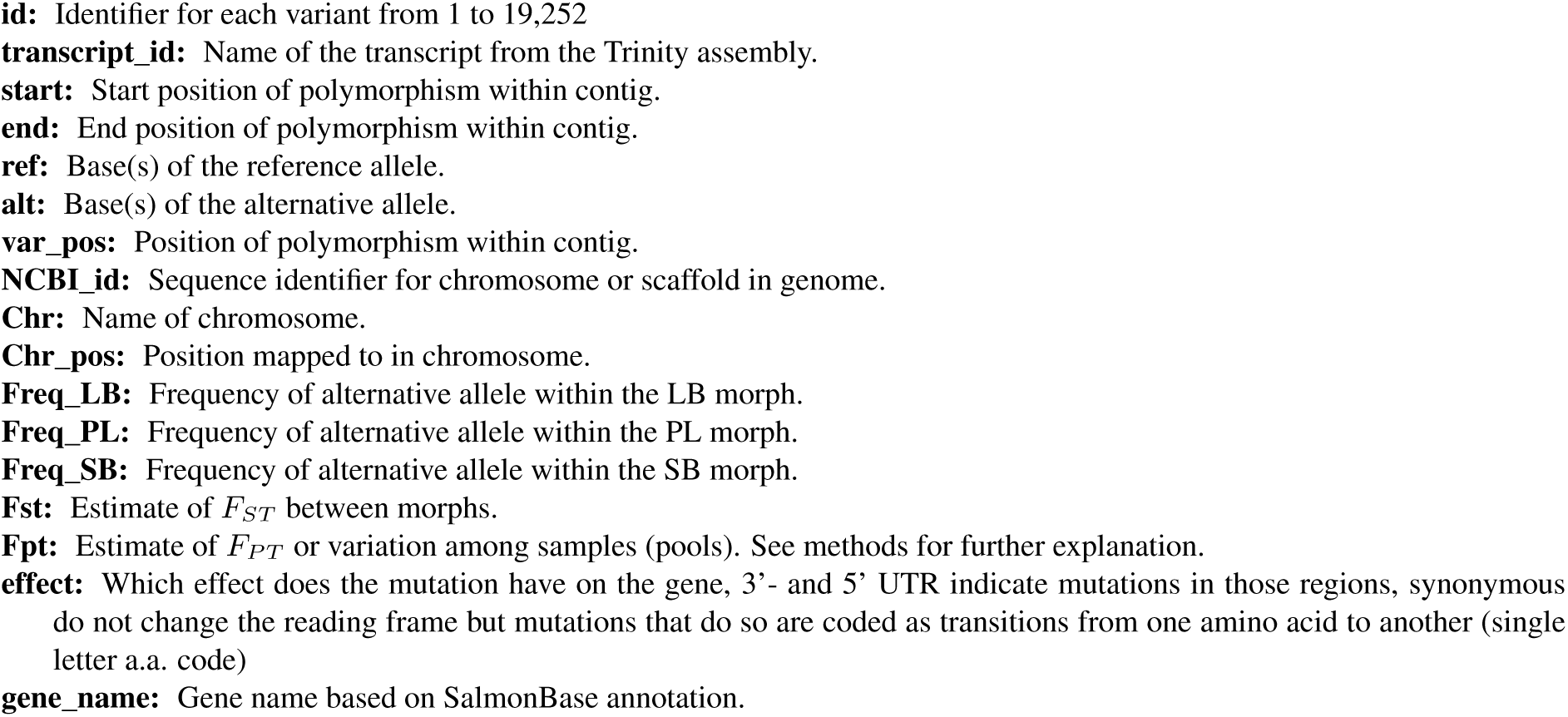
Tab-delimited text file with the position, alleles, alternative frequencies within morphs, F-statistics between morphs and samples and predicted effect of the variant on protein composition for all the variants after the final filtering step. On figshare: doi:10.6084/m9.figshare.8705888

**Table S2:** VCF-file with the final set of variants after all filtering steps.

On figshare: doi:10.6084/m9.figshare.8718695

**Table S3:**
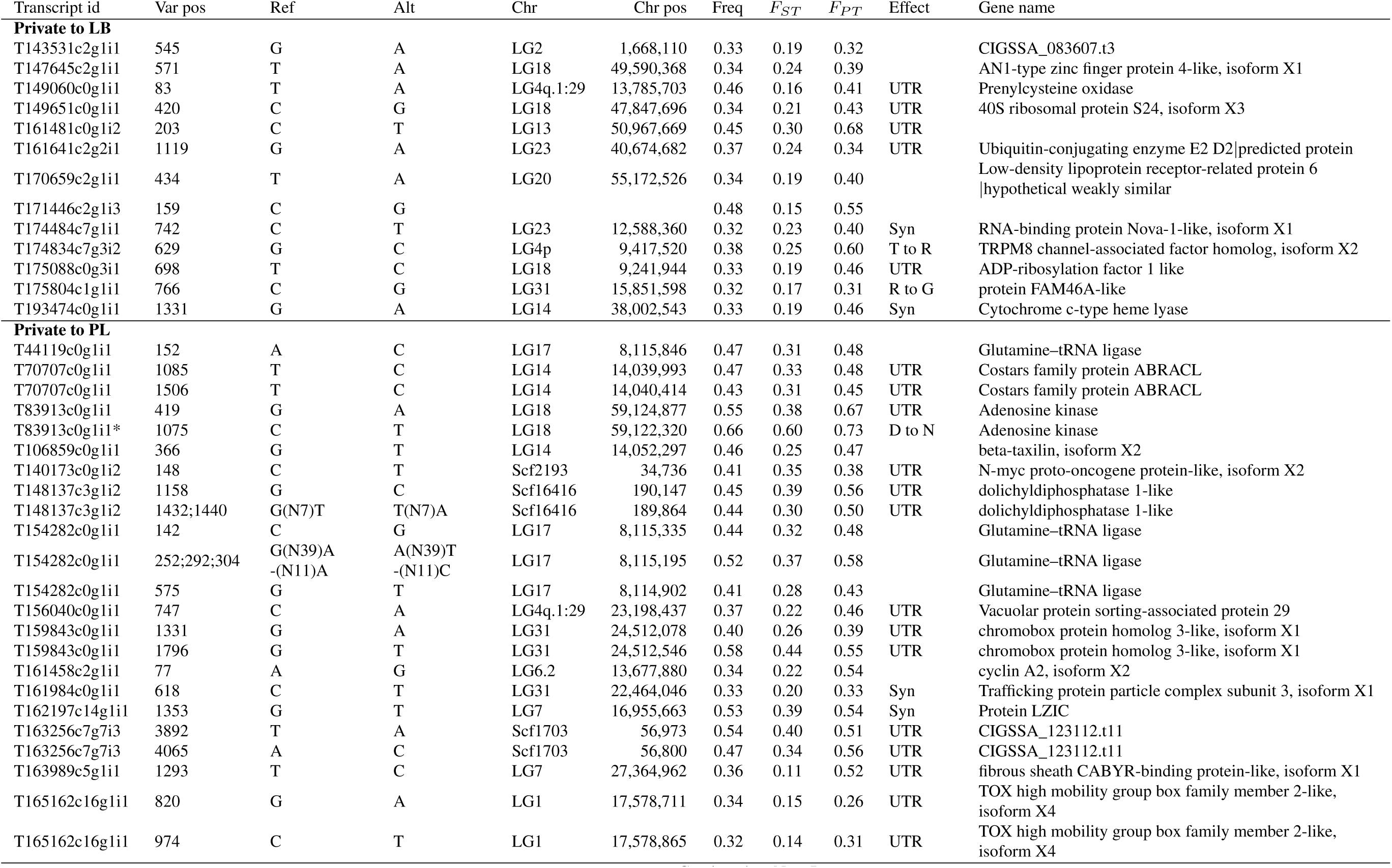
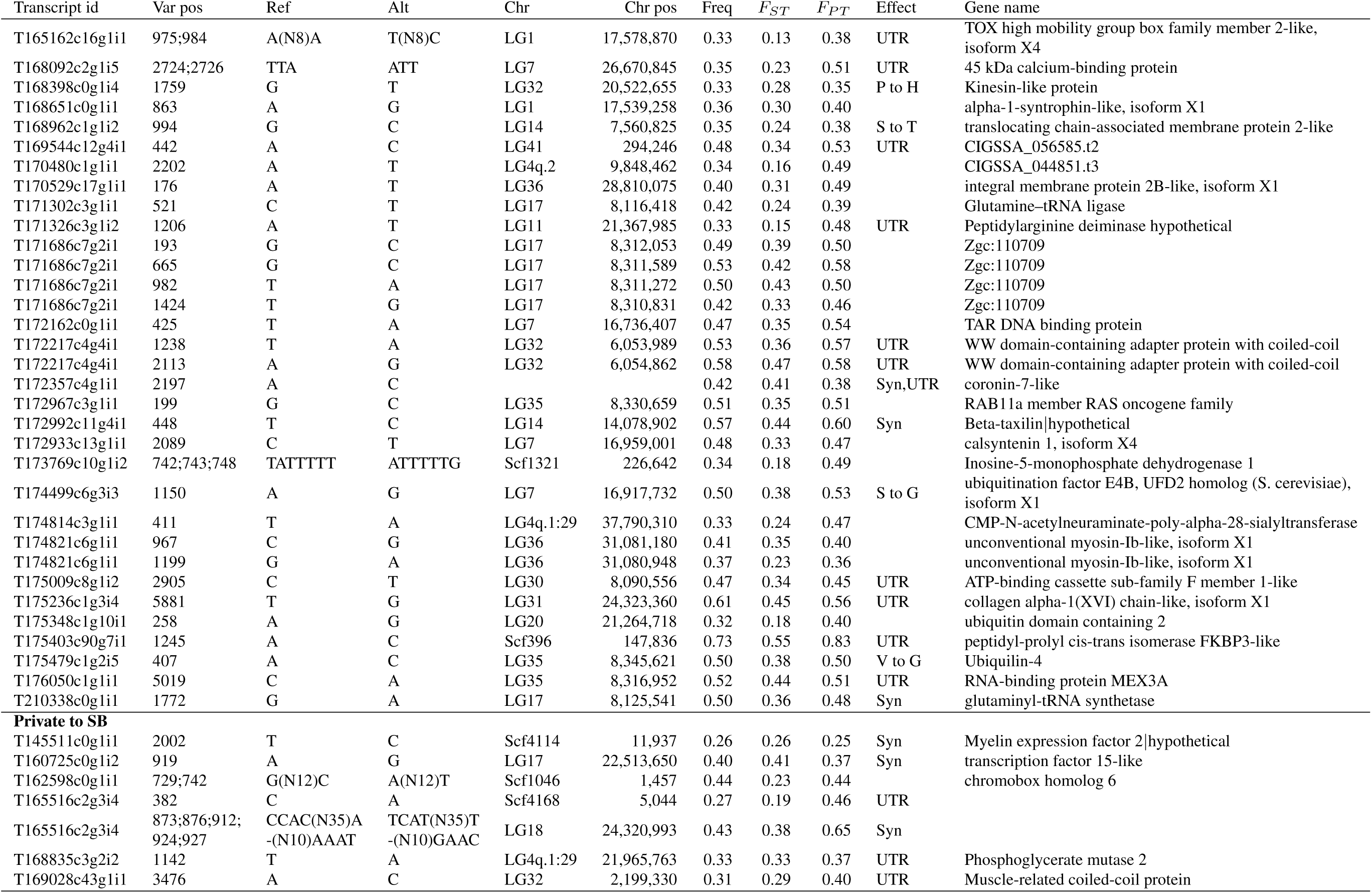
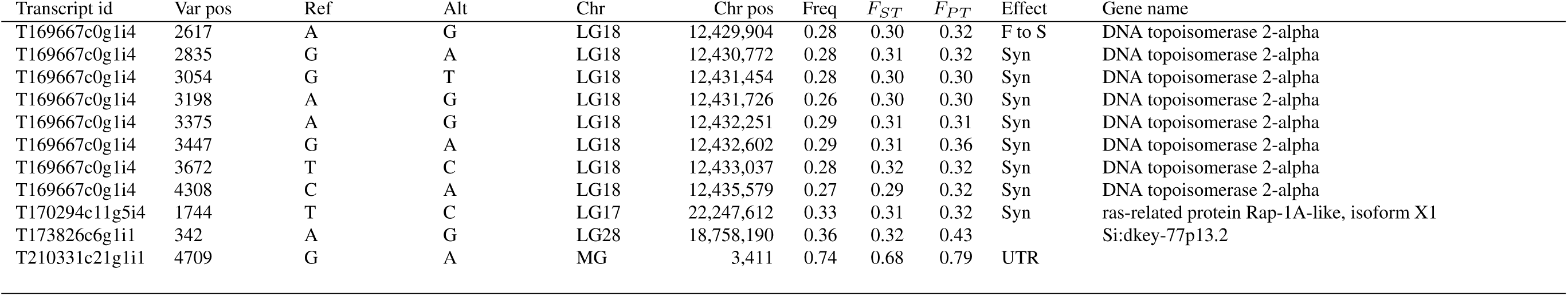
Variants only found within one morph. Position within contig and genome, alleles, frequency within the morph, *F*_*ST*_-values and predicted effect are shown. *Validated in KASP-assay

**Table S4:**
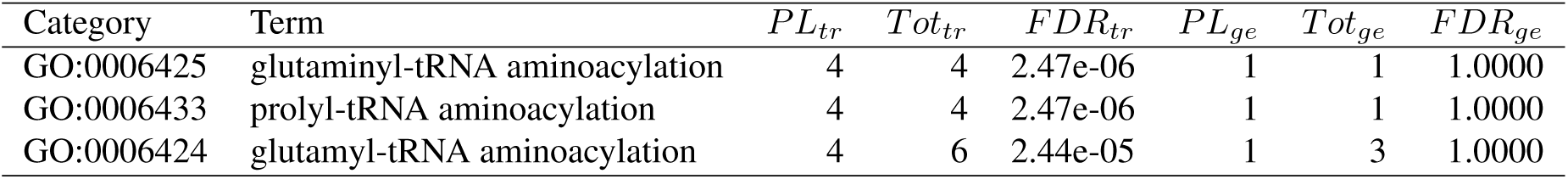
Gene ontology categories enriched in variants private to PL-charr (*p <* 0.01 in the other morphs) on the transcripts (*tr*) and gene (*ge*) level. The number of transcripts and genes with observed private variants (*PL*_*tr*_ and *PL*_*ge*_) and the total number of transcripts and genes tested in the category (*T ot*_*tr*_ and *Tot*_*ge*_) are shown. The multiple testing corrected p-value or false discovery rate (*FDR*) is also shown for both levels.

**Table S5:**
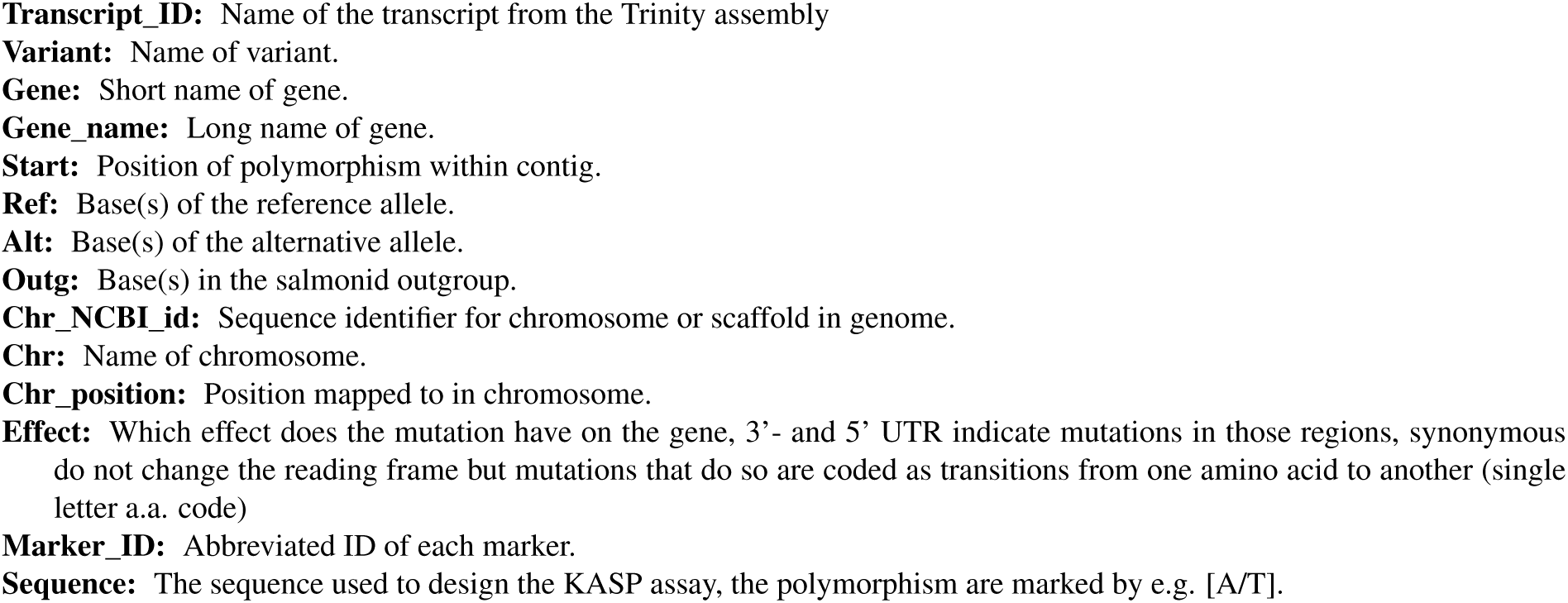
Genetic polymorphisms studied in the population sample of Arctic charr. On figshare: doi:10.6084/m9.figshare.8719784

**Table S6:** Datafile of the genotypes for the 22 markers scored in the population samples from the four sympatric charr morphs (coded by bases, missing data indicated by “NA”).

On figshare: doi:10.6084/m9.figshare.8721317

**Table S7:**
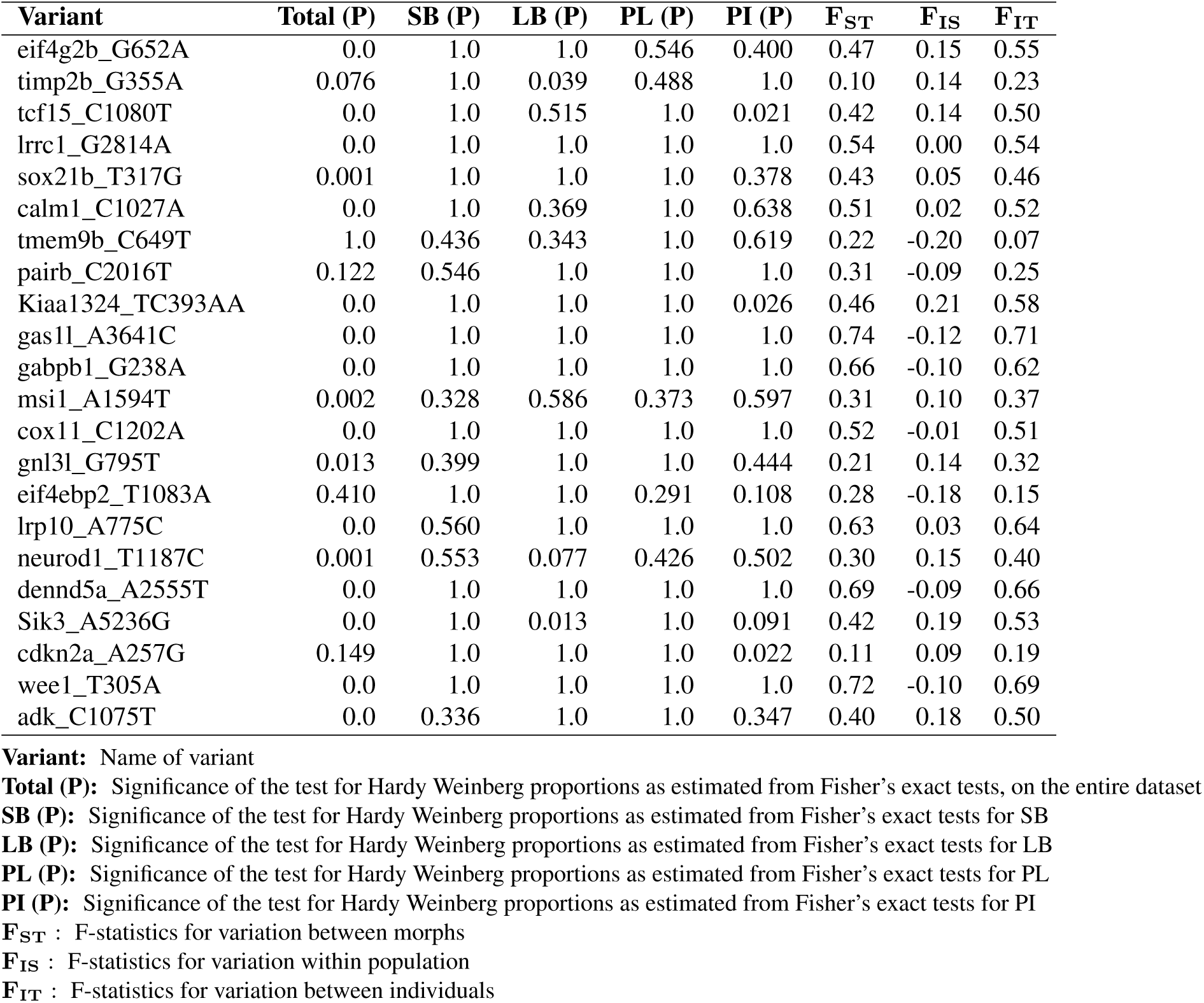
Estimates of F-statistics for the entire KASP data and tests of Hardy-Weinberg proportions for the entire KASP dataset (Total) and individual morphs.

**Table S8:** Datafile of the LD (*r*^2^) for all pairs of markers, by morph On figshare: doi:10.6084/m9.figshare.8721455

On figshare: doi:10.6084/m9.figshare.8721455

## Supplementary Figures

**Figure S1:**
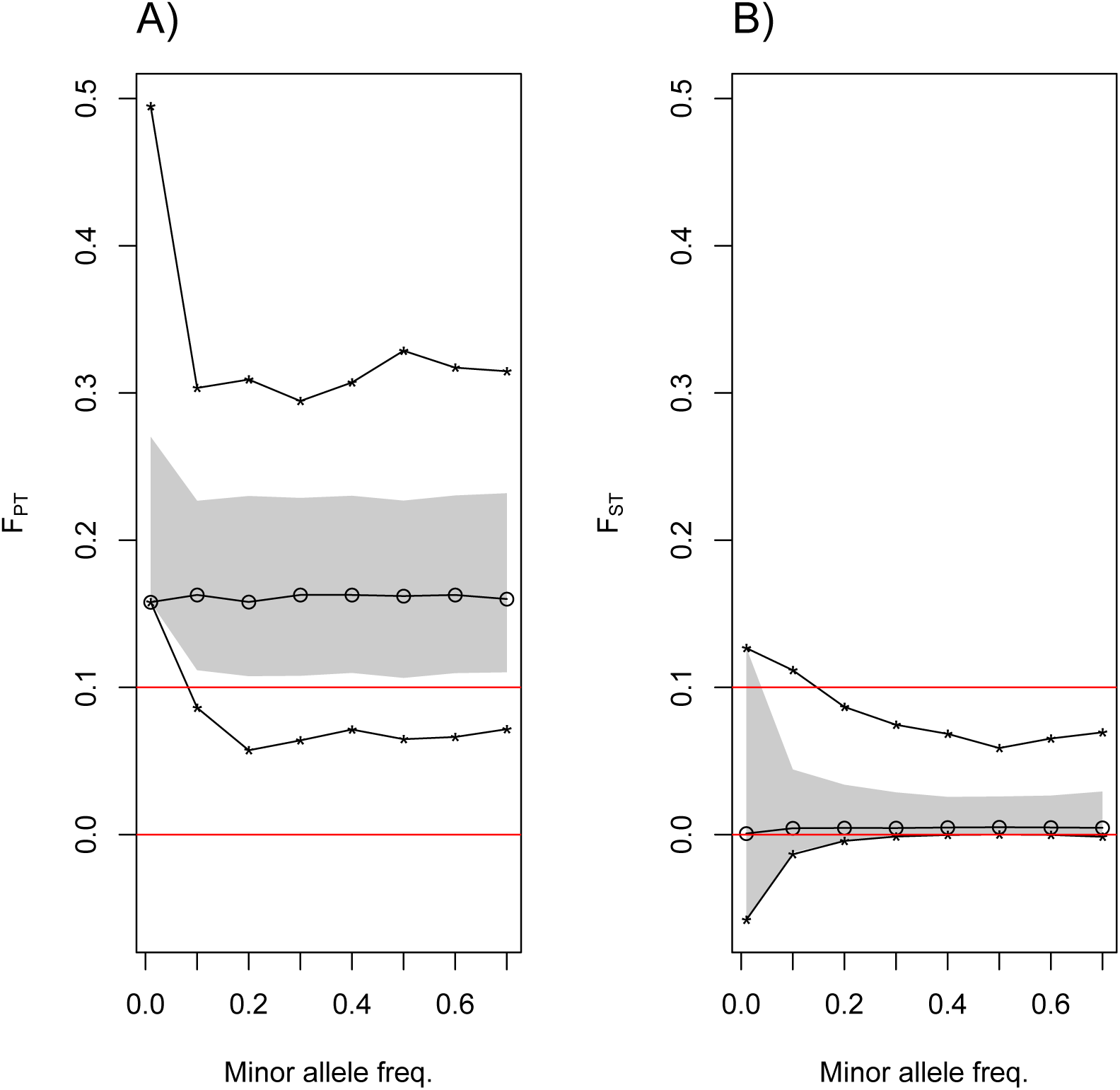
Results from simulations on F-statistics for a biallelic variant with the same allele frequency among morphs. The gray area indicate the 95% confidence area. The median, minimum and maximum values are shown by dots and lines. A) Shows *F*_*PT*_ values and B) *F*_*ST*_ values. As expected *F*_*PT*_ values are high and we chose 0.1 (red horizontal line) as cutoff as it is outside the 95% confidence area in the simulations.

**Figure S2:**
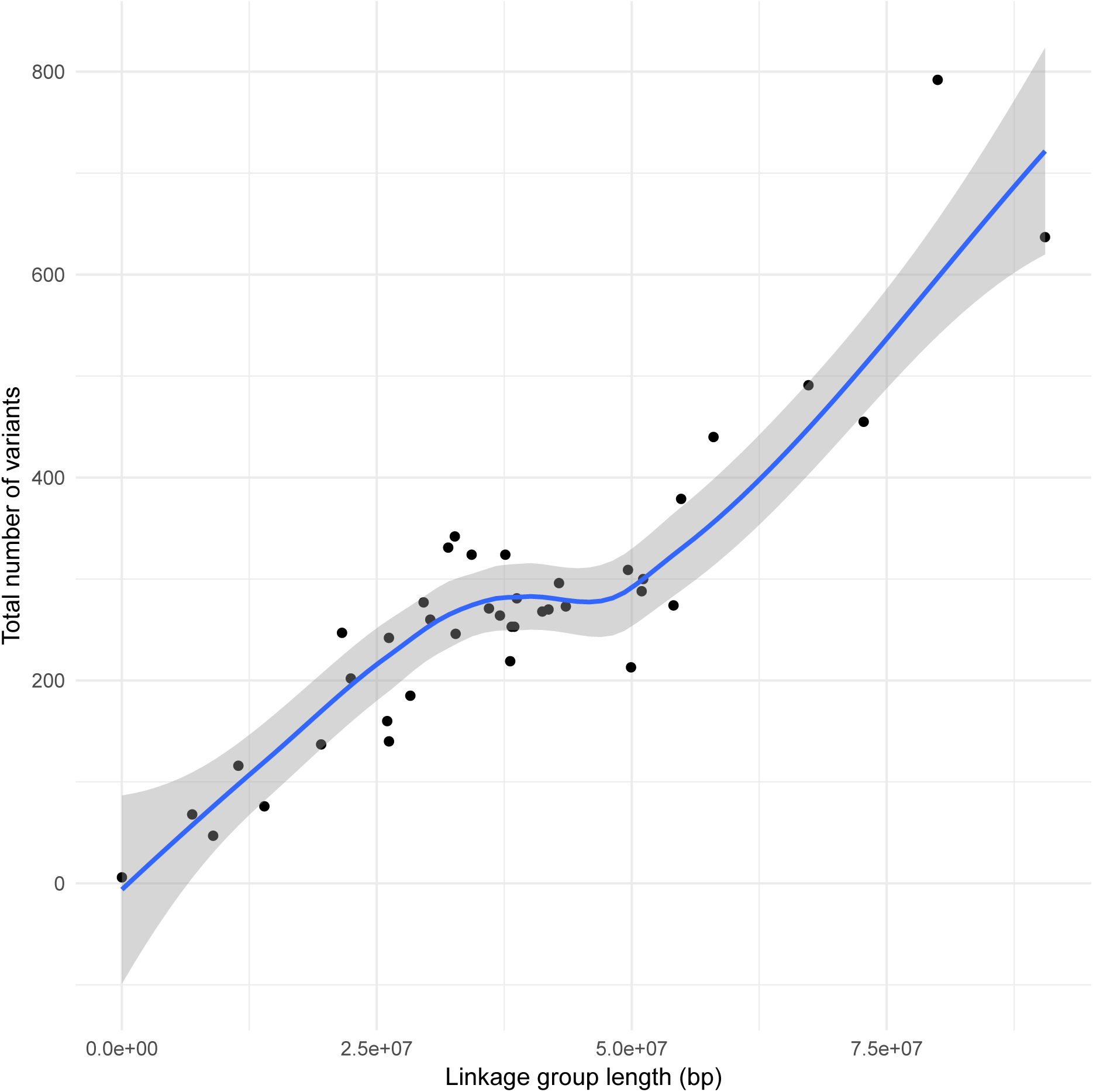
Total number of transcriptome variants by linkage group length. A loess smooth curve (blue) with 95% confident interval (gray) is also graphed.

**Figure S3:**
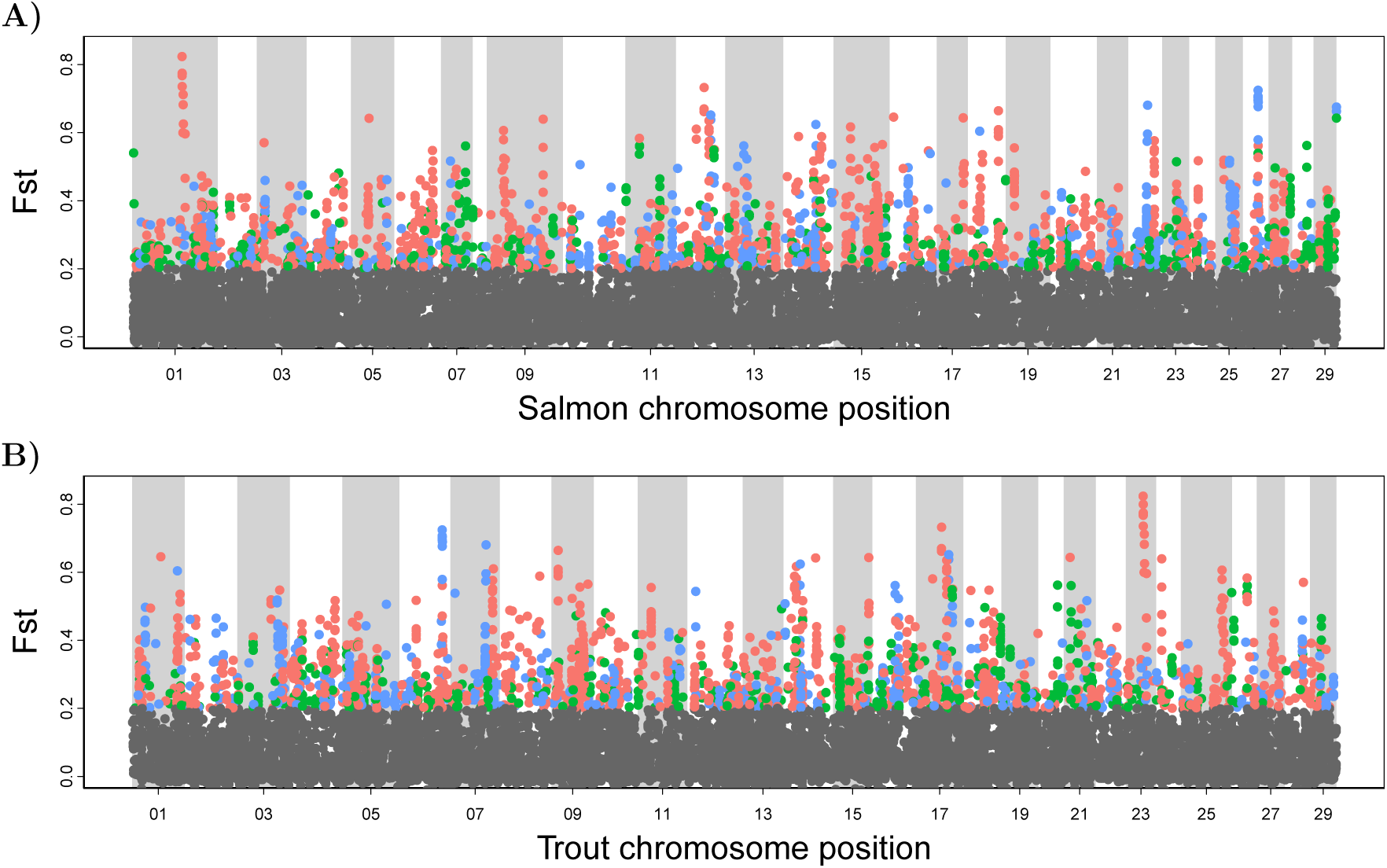
*F*_*ST*_ values plotted by position of variants on the salmon **A)** and rainbow trout **B)** genome. The colors indicate which morphs differs most strongly in allele frequency from the other two for variants with *F*_*ST*_ *>* 0.2. Red PL, blue SB and green LB

**Figure S4:**
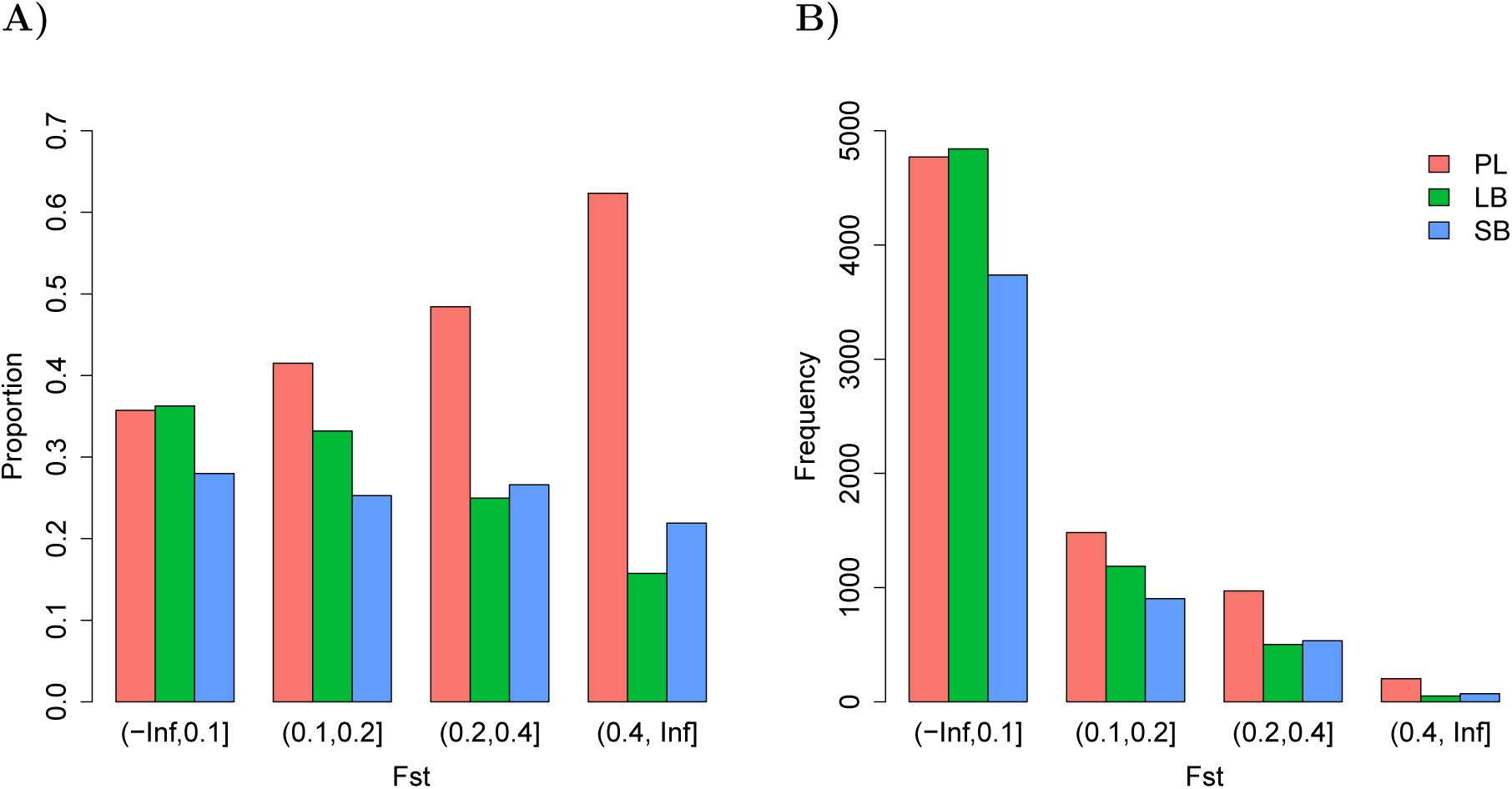
**A)** Proportion of variants (within *F*_*ST*_ groups) for different *F*_ST_-values grouped by the morph with highest deviation in allele frequency from the other two. **B)** The number of variants in each *F*_*ST*_ category grouped by the morph with highest deviation in allele frequency from the other two. The legend in **B)** also applies to **A)**.

**Figure S5:**
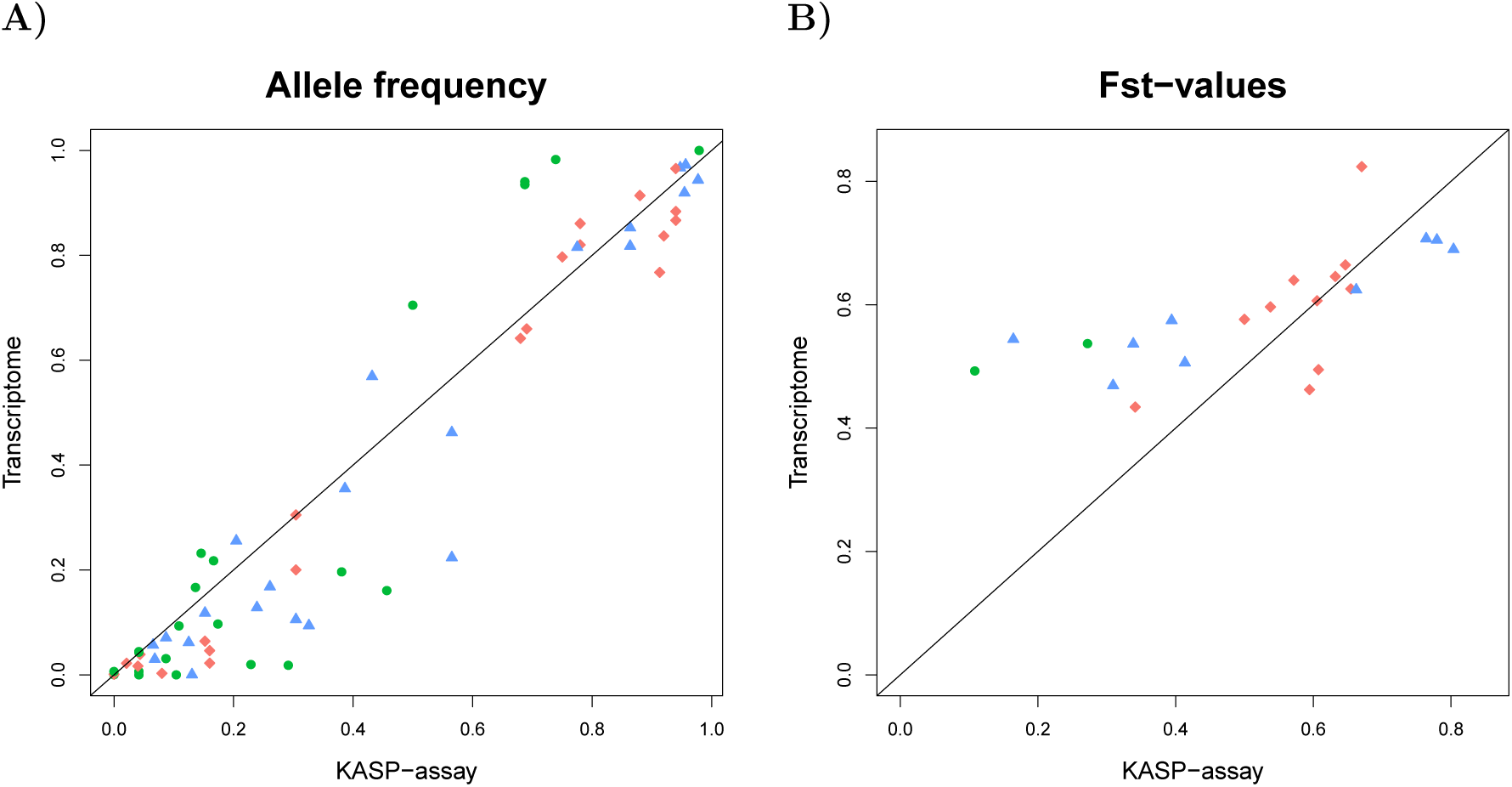
Tight relationship was found between the all1ele frequencies and *F*_ST_ values estimated from the transcriptome and the population genetic sample (Kasp assay). **A)** The allele frequencies for the 22 markers were estimated for each morph (color coded) and the estimates from the two methods show high positive correlation (Kendall’s *τ* = 0.76, Pearson’s *r* = 0.96, *p <* 0.0001). The diagonal line represents the 1:1 relationship. **B)** *F*_*ST*_-values calculated for the transcriptome and the population sample had comparatively weaker association (Kendall’s *τ* = 0.58, Pearsons’s *r* = 0.70, *p <* 0.001). Notably, two markers deviated substantially between the two methods (in *timp2b* and *cdkn2a*), with higher *F*_*ST*_ in the transcriptome compared to the population sample (due to underestimation of some rarer allele frequencies in the transcriptome). The colors indicate which morphs shows the highest deviation from the other two in mean allele frequency for each marker in the transcriptome (SB: blue, LB: green and PL: red).

**Figure S6:**
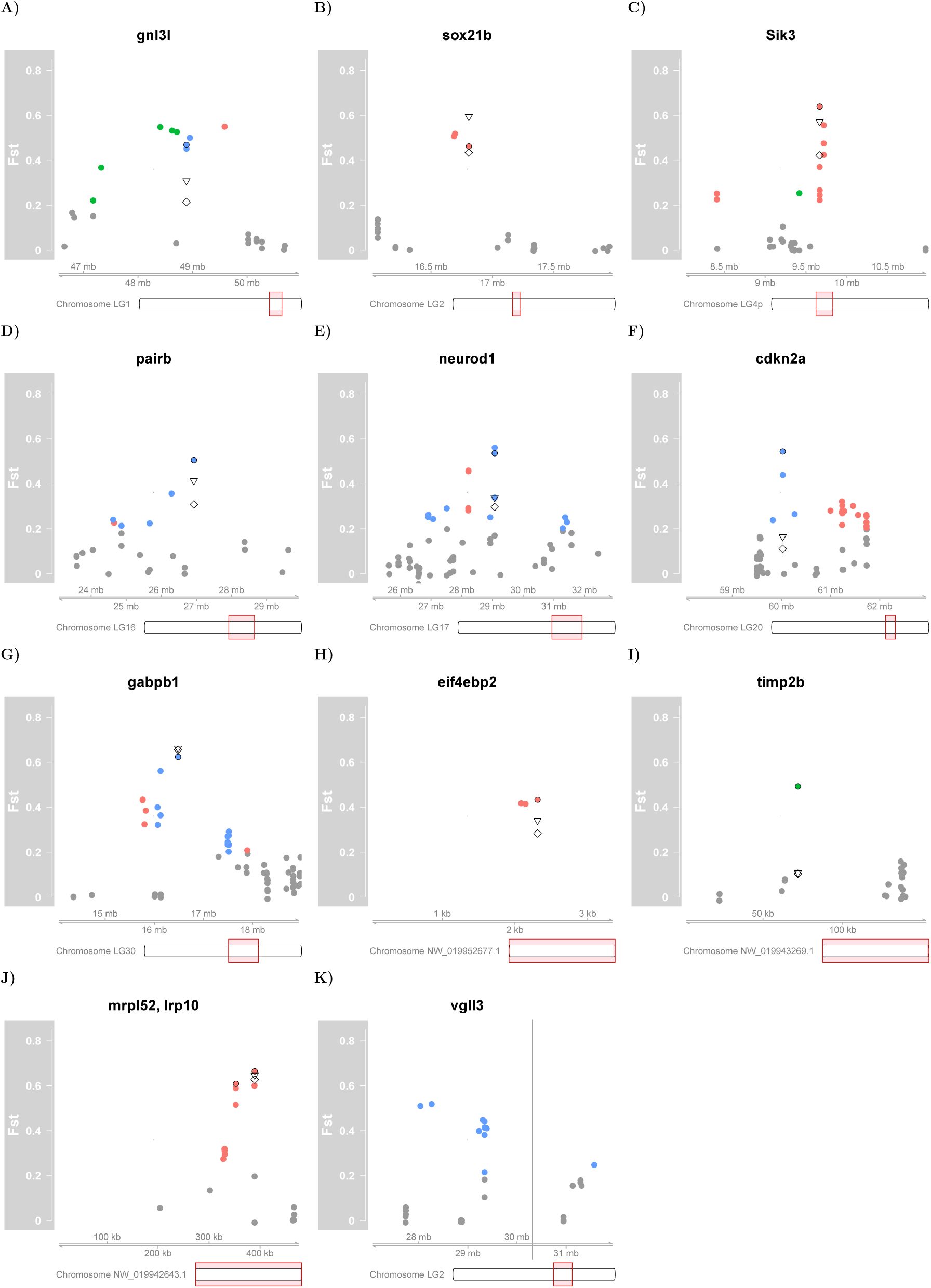
**A-J)** Detailed view of chromosomal regions of variants in KASP-assay and nearby transcriptome variants (not-validated). The colored dots mark *F*_*ST*_ values from the transcriptome as in Fig. 3 and the variants taken for validation are marked with a filled black circl (∘). The triangles (∇) show the *F*_*ST*_ value from the KASP-assay for the three transcriptome morphs (PL, SB and LB) and the diamonds (⋄) *F*_*ST*_ for all morphs (including PI). **K)** Detailed view of variants nearby the *vgll3* locus. The vertical line represents the location of the *vgll3* gene, not transcribed and thus no variants detected, using the same color code (SB: blue, LB: green and PL: red) for variants. 38

## Notes

#### Summary of Updates

In this revised manuscript we trimmed the text, tried to provide a clearer focus for the study, modifying introduction, results and discussion and have taken into account the very helpful comments provided by the reviewers. For instance, we have added more details regarding generation of crosses and sampling and discuss them in the results. We also disclose that two errors were discovered in the genotyping-assay, that were corrected as follows: 1) A individual labeled as an SB-charr in previous version, was actually a PL-charr (based on a photograph). This was the stray SB that previously grouped with the PL-charr. 2) The peculiar "result" of strong LD between cdkn2a and gnl3l which mapped to different chromosomes was due to technical error. These two markers had been run at the same time and there was a primer mixup. Both primers were rerun that gave the same results for cdkn2a but quit different for gnl3l. These two errors led us to rewrite the results on genotyping (KASP-assay) and redo associated figures and tables. But these corrections did not affect the general conclusions, rather strengthened the correlation of allele frequencies in the transcriptome and the KASP data. Finally, we changed the format of references to a Harvard style and uploaded supplementary tables and files on figshare.

https://doi.org/10.6084/m9.figshare.c.4565735.v1

